# Sox11 is enriched in myogenic progenitors but dispensable for development and regeneration of skeletal muscle

**DOI:** 10.1101/2023.03.30.534956

**Authors:** Stephanie N. Oprescu, Nick Baumann, Xiyue Chen, Qiang Sun, Yu Zhao, Feng Yue, Huating Wang, Shihuan Kuang

## Abstract

Transcription factors (TFs) play key roles in regulating the differentiation and function of stem cells, including muscle satellite cells (MuSCs), a resident stem cell population responsible for postnatal regeneration of the skeletal muscle. Sox11 belongs to the Sry-related HMG-box (SOX) family of TFs that play diverse roles in stem cell behavior and tissue specification. Analysis of single-cell RNA-sequencing (scRNA-seq) datasets identify a specific enrichment of *Sox11* mRNA in differentiating but not quiescent MuSCs. Consistent with the scRNA-seq data, *Sox11* levels increase during differentiation of murine primary myoblasts in vitro. scRNA-seq data comparing muscle regeneration in young and old mice further demonstrate that *Sox11* expression is reduced in aged MuSCs. Age-related decline of *Sox11* expression is associated with reduced chromatin contacts within the topologically associated domains. Unexpectedly, Myod1^Cre^-driven deletion of *Sox11* in embryonic myoblasts has no effects on muscle development and growth, resulting in apparently healthy muscles that regenerate normally. Pax7^CreER^ or Rosa26^CreER^ driven (MuSC-specific or global) deletion of *Sox11* in adult mice similarly has no effects on MuSC differentiation or muscle regeneration. These results identify Sox11 as a novel myogenic differentiation marker with reduced expression in quiescent and aged MuSCs, but the specific function of Sox11 in myogenesis remain to be elucidated.

## Background

Skeletal muscle makes up nearly 40% of the total body mass and is critical for balance, movement and maintaining quality of life (1). As a surface tissue, the skeletal muscle is prone to various injuries. Mammalian skeletal muscles harbor a resident population of adult stem cells, known as muscle satellite cells (MuSCs), that in response to external stimuli such as an injury, activate, proliferate, and differentiate to repair the injured muscle. While various infiltrating and resident cells are necessary for clearing debris, remodeling the extracellular matrix, and modulating the regenerative environment, MuSCs critically contribute to the repair by fusing together to generate new fibers to restore muscle function (2). To functionally repair the muscle, MuSCs must first exit their quiescent state, proliferate to expand the pool, and then either commit to the myogenic program and differentiate or maintain their stem-like state and self-renew to replenish the adult MuSC pool. This balance of self-renewal and differentiation is critical for muscle function, and dysregulation can lead to exhaustion of the MuSC pool and impair subsequent regenerative capacity under pathological conditions (3, 4, 5). While external signals in the environment impact the MuSC response, these signals must be integrated within the cell and ultimately affect cell identity and cell fate decisions (6). How a cell responds to various cues is mediated by its transcriptional state, underscoring the importance of defining key transcriptional regulators in cell identity and cell states.

Muscle stem cell activation, proliferation, and differentiation during muscle repair recapitulates various aspects of the myogenic developmental process, and the major transcriptional regulators of this process are well understood (6, 7, 8, 9, 10, 11, 12, 13). The best-known transcription factors are Pax3/Pax7 and the myogenic regulatory factors (MRFs, which include Myf5, Myf6, Myod1, and Myogenin) (6, 7, 8, 9, 14, 15, 16, 17, 18). The expression of these factors is MuSC-state specific, with Pax3/Pax7 primarily expressed in quiescent and proliferating MuSCs, while the MRFs are differentially expressed along the myogenic lineage (19). Cells expressing Pax7 are generally refractory to differentiation cues and will maintain their self-renewing state via controlling transcription of quiescence-related genes, mainly influenced by external signals but also regulated by the transcriptional state of MuSCs (20, 21). Thus, transcriptional states and integration of external environmental signals is relatively complex. Notch signaling is one of the best characterized pathways that plays a critical role in maintaining the balance of MuSC quiescence and self-renewal (20), through regulation of Hes/Hey family transcriptional factors. During development, the Notch ligand Delta-like 1 (Dll1) regulates myogenic differentiation and maintenance of progenitors, as high expression of the Notch intracellular domain (NICD) supports their proliferation (22, 23). Conversely, in adult MuSCs high NICD expression maintains their quiescence, in part by targeting the expression of niche-related collagen genes and miR-708, which limits migration and further reinforces quiescence through niche interactions (24, 25, 26). As MuSCs proliferate, high expression NICD supports *Pax7* expression to promote the self-renewal of MuSCs (27, 28). Interestingly, changes in the expression of Notch ligands and reduced p53 expression have been linked to age-related functional decline of MuSCs (29, 30). In addition to the critical role of Notch signaling, Wnt signaling has also been shown to mediate MuSC function, also through altering a transcriptional program mediated by β-catenin. For example, Wnt1+ cells preferentially activate *Myf5*, while cells expressing Wnt7a preferentially activate *Myod* (31, 32). In response to muscle injury, canonical Wnt/ β-catenin signaling promotes commitment of myoblasts by regulating the expression of follistatin (33). Age-related increase in Wnt signaling was also found to mediate the conversion of MuSCs to a more fibrogenic-like state, leading to impaired regeneration (34). Overall, these findings exemplify the role of transcriptional factors in mediating external signals to regulate MuSC state and function.

To identify factors that regulate transcriptional states and may support or mediate MuSC fate transitions, we employed published and newly generated single-cell RNA-sequencing data to identify *Sox11* as a transcriptional factor enriched in differentiating MuSCs. The SOX protein family is best known for their role in development, embryonic stem cells, tissue specification and sex determination (35, 36). Among the 20 SOX proteins, Sox11 is a member of the SoxC sub-family, which also includes Sox4 and Sox12, all of which are widely expressed during embryogenesis and are developmentally required (Fig. 1A) (36, 37). Although Sox4 and Sox11 exhibit some redundancy, they are both critical for cell and embryo survival. *Sox11-/-* mice are born viable but do not survive past 24 hours due to impaired organogenesis such as under-mineralized bones and heart malformations (38, 39, 40). Sox11 is broadly required for survival of neural and mesenchymal progenitors and mediates the proliferation of neural progenitors in the central nervous system, while promoting precursor differentiation in the peripheral nervous system (7, 38, 41, 42, 43). These findings underscore the complex, cell, and tissue-type specific regulatory role for Sox11 during development.

**Figure 1.**
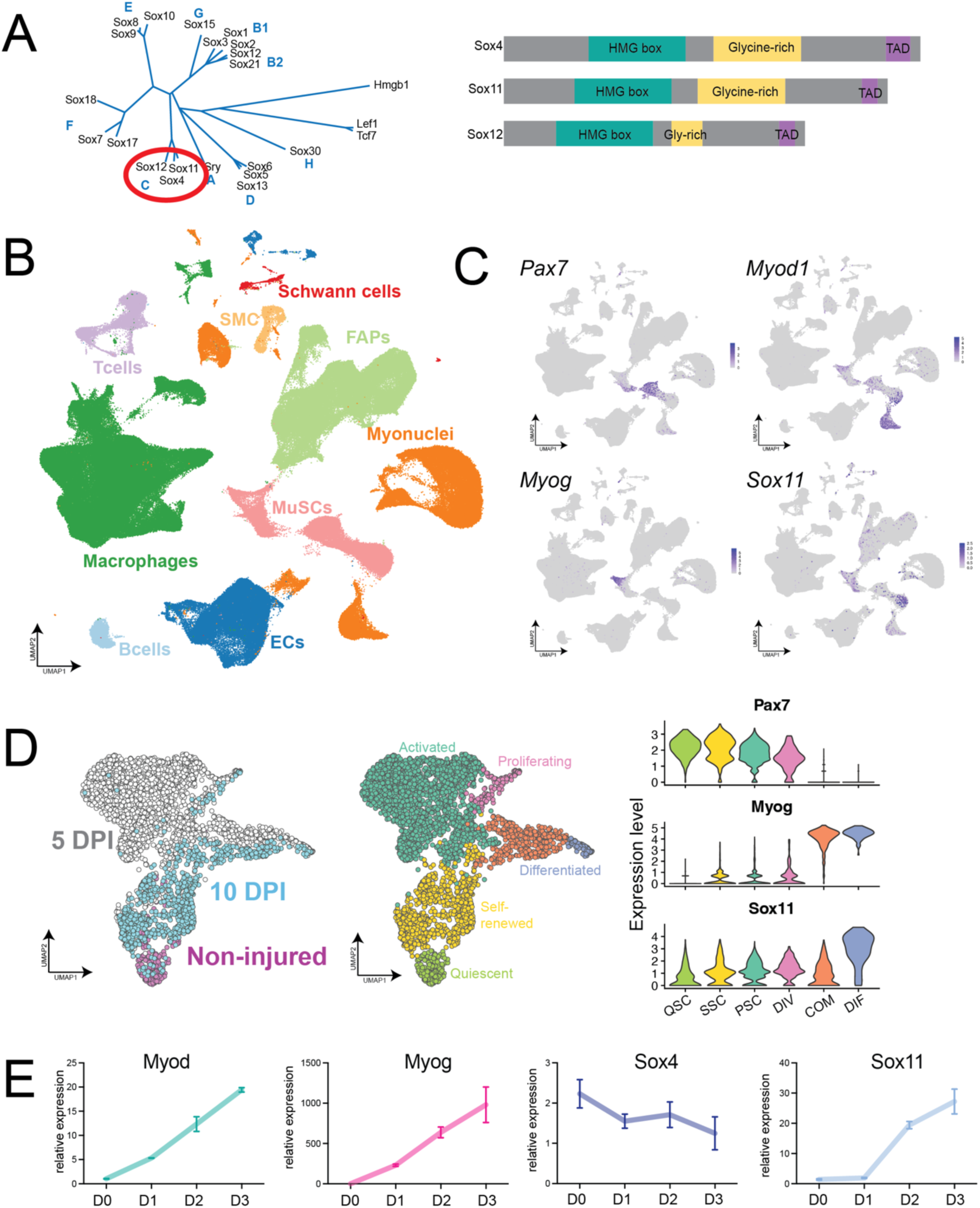
Single-cell RNA-sequencing identified *Sox11* expression in differentiating muscle stem cells. **A.** Phylogenetic neighbor-joining tree for the high-mobility group (HMG) domain containing Sox factors highlighting the SoxC sub-family and their domain phylogenetic similarity for mammals (89). **B.** UMAP plot of McKellar et al., aggregated dataset (GEO accession: GSE162172), broadly classified by cell type. **C.** UMAP-based gene expression plots from McKellar dataset for *Pax7*, *Myod1*, *Myog* and *Sox11*. **D.** UMAP projection of scRNA-seq of MuSCs sorted from non-injured, 5 and 10 DPI muscle, colored by time-point and cluster identity (left panel), (GEO accession: GSE150366). Violin plot of *Pax7*, *Myog* and *Sox11* based on scRNA-seq dataset to highlight expression in different MuSCs clusters. **E.** qRT-PCR on RNA isolated from proliferating (D0) myoblasts and myoblasts induced to differentiate for 1, 2 and 3 days to detect the changes in expression of *Myod1*, *Myogenin*, *Sox11* and *Sox4*.

The role of Sox11 during adulthood has also been investigated in several disease and regeneration settings. High expression of *Sox11* in various cancers is generally correlated with poor prognosis as it appears to support the epithelial to mesenchymal transition of cancer cells (44). Studies on the role of Sox11 in tumorigenesis indicate that Sox11 regulates genes involved in Wnt signaling and the Notch pathway, both of which are imperative for MuSC function (27, 28, 33, 34, 45, 46, 47, 48). Sox11 also regulates *Tead2* expression to support the survival and proliferation of osteoblasts and mesenchymal cells while regulating *Osterix* and *Runx2* to promote osteoblast and mesenchymal cell differentiation (49). Interestingly, Sox11 was also shown to mediate sensory nerve regeneration, as knock-down of *Sox11* RNA inhibited regeneration *in vivo* in adult mice (41, 50). Additionally, Sox11 (and another member of the SoxC subfamily, Sox4) was found to reactivate embryonic developmental programs to support skin wound repair by inhibiting premature differentiation (51). However, the expression and function of Sox11 in MuSCs and muscle regeneration have not been investigated.

Several Sox proteins have been reported to play a role in myogenesis. For example, Sox6 functions to repress slow-fiber type gene expression, while Sox8 negatively regulates MuSC differentiation (52, 53, 54). Additionally, a previous study identified another member of the SoxC sub-family, Sox4, is important for differentiation of the muscle cell line, C2C12, by targeting the *Cald1* promoter (55). However, unlike *Sox11*, we found that *Sox4* expression is not restricted to MuSCs and does not increase during primary myoblast differentiation, consistent with a previous study that detected increased expression of *Sox11* during myoblast differentiation (52). Given the enrichment of Sox11 in myogenic cells and the broad regulatory role of Sox11 in other non-muscle tissues, we hypothesized that Sox11 may play a unique role in myogenesis. In the present study, we used multiple scRNA-seq datasets from acutely injured skeletal muscle to identify enriched *Sox11* expression in differentiating MuSCs. We subsequently investigated the requirement of Sox11 muscle progenitor function using various conditional knockout mouse models. While these data suggest that Sox11 is dispensable for normal MuSC function under the evaluated conditions, it adds to our understanding of the potential redundancy of the SOX family members in myogenesis.

## Methods

### Muscle injury and sample processing for single-cell RNA-sequencing

Hindlimb muscle of 3 *Pax7*^nGFP/+^ male mice at 3-4 months of age (young) and >20 months of age (old) was injured via intramuscular injection of 50 µL 10µM cardiotoxin. Hindlimb muscles were dissected at 7 DPI, digested to release mononuclear cells, and sorted via fluorescence activated cell sorting to select for live, single cells as previously described (56, 57).

### Single-cell RNA-sequencing

scRNA-sequencing was performed using the 10X Genomics 3’ v2 kit, following their protocol targeting recovery of 10,000 cells. Libraries were constructed per the manufacturer’s instructions, sequenced on Illumina’s NovaSeq platform. Reads were aligned to the mouse genome mm10/Grcm38 using the CellRanger 2.1.0 software and additional analysis was performed in R.

### Quality control, dimensionality reduction, and visualization

Seurat 3.1.0 in R was used to analyze CellRanger output following a broadly similar pipeline as previously described (58). Both young and old samples were merged and filtered for cells with less than 15% reads mapping to mitochondrial genes, gene counts less than 6,000 per cell and no more than 60,000 reads. Both Seurat’s SCTransform function and log-transformation methods were used to normalize and scale the data and subsequently compare results, which yielded similar outcomes (data not shown) (59). Dimensionality reduction was performed through Principal Component Analysis (PCA) and top principal components were selected by evaluating elbow plots. Clustering and UMAP embedding parameters were based on the top 10 PCs and embedded in 2-dimensions for visualization. FindAllMarkers() function was used to identify gene enriched in each cluster, which were used to manually label cell types. All genes considered for cell-type classification had a *P-*value of less than 0.0001 using a Mann-Whitney Wilcoxon test. To perform the sub-clustering, we used Seurat’s subset function to extract the cell types of interest (MuSCs), we extracted the raw RNA counts for each assayed cell type to subset, re-scaled the data using the SCTransform function, performed dimensionality reduction, clustering and UMAP visualization. We then compared gene expression based on cell sample (i.e., age). Additional scRNA-sequencing data is based on previously analyzed and published datasets (60, 61).

### Gene expression analysis of RNA-seq data

The raw reads of total RNA-seq were processed following the procedures described in previous publication. Briefly, the adapter and low-quality sequences were trimmed and the reads shorter than 50 bp were discarded. The clean reads were mapped to mouse genome (mm9) with Bowtie2 (V 2.1.1). Cufflinks (V 2.2.1) were then used to estimate gene expression level in Fragments Per Kilobase per Million (FPKM). Genes were annotated as differentially expressed if the change of expression level is greater than 2 folds between two stages/conditions.

### ChIP-seq data analysis

Raw ChIP-seq reads were processed as previously described(62). Briefly, the adapter and low-quality sequences were trimmed from 3’ to 5’ ends by Trimmomatic (V 0.36) and the reads shorter than 36 bp were discarded. Subsequently, the preprocessed reads were aligned to the mouse genome (mm9) using Bowtie2 (v2.3.3.1). The duplicate reads were removed by Picard (http://broadinstitute.github.io/picard). Peaks were then identified by MACS2 (V 2.2.4) with q-value equal to 0.01 by using the IgG control sample as background.

### *In situ* Hi-C data processing

The *in-situ* Hi-C data was processed with HiC-Pro (v2.10.0)(63). First, adaptor sequences and poor-quality reads were removed using Trimmomatic (ILLUMINACLIP: TruSeq3-PE-2.fa:2:30:10; SLIDINGWINDOW: 4:15; MINLEN:50). The filtered reads were then aligned to reference genome (mm9). All aligned reads were then merged together and assigned to restriction fragment, while low quality (MAPQ<30) or multiple alignment reads were discarded. Invalid fragments including unpaired fragments (singleton), juxtaposed fragments (re-legation pairs), un-ligated fragments (dangling end), self-circularized fragments (self-cycle), and PCR duplicates were removed from each biological replicate. The remaining validate pairs from all replicates of each stage were then merged, followed by read depth normalization using HOMER (http://homer.ucsd.edu/homer/interactions/HiCpca.html) and matrix balancing using iterative correction and eigenvector decomposition (ICE) normalization to obtain comparable interaction matrix between different stages.

### Identification and analysis of TADs

Normalized contact matrix at 10 kb resolution of each time point was used for TAD identification using TopDom (v0.0.2)(64). In brief, for each 10-kb bin across the genome, a signal of the average interaction frequency of all pairs of genome regions within a distinct window centered on this bin was calculated, thus TAD boundary was identified with local minimal signal within certain window. The insulation score of the identified TAD border was also defined as previously described, which used the local maximum on the outside of TAD to minus the local minimum on the inside of TAD of each boundary bin.

### scTenifoldKnk *in silico* knock-out analysis

Functional analysis of *Sox11* was conducted using the R package of *scTenifoldKnk* (65). A single-cell gene regulatory network (scGRN) was conducted on muscle satellite cells from the control samples in our previously published scRNA-seq dataset ((61); GSE150366). Then, the expression of *Sox11* was set to zero from the constructed scGRN to build their own corresponding “pseudo-knockout” scGRN. Perturbed genes by this virtual knockout were quantified by comparison of the “pseudo-knockout” scGRN to the original scGRN. Those significantly affected genes were used for functional enrichment analysis (GO and KEGG) to show changes in biological processes caused by in silico knockout.

### Animals

Animals used in this study were: Pax7^CreERT2^(#017763), Rosa26^CreER^ (#008463), Myod1^Cre^ (#014140) were purchased from Jackson Laboratory as the respective stock number. Sox11^flox/flox^ were a gift from Dr. Veronique Lefebvre (Children’s Hospital of Philadelphia). Mouse genotypes were evaluated by genomic DNA isolation from the ear and determined via polymerase chain reaction (PCR). For mice purchased from Jackson laboratory, primers and protocols published with each. For Sox11, genotyping for the floxed allele was determined via PCR as previously described (66). Recombination PCR and recombination PCR was determined using primers specific for the recombined allele on DNA isolated from whole muscle or myoblasts treated with MeOH or 4OH, using a previously described protocol (66). 3–6-month-old mice were used, were always age and litter matched. Littermate controls included both Cre-positive heterozygous floxed mice and Cre-negative mice. No sex-specific differences were observed. All procedures and mice were approved and housed according to the Purdue Animal Care and Use Committee standards.

### Tamoxifen

For *Pax7*^CreER^ and *Rosa26*^CreER^ mice, tamoxifen was in to induce recombination of the floxed allele. Tamoxifen (100mg/mL) was administered via intraperitoneal injection of 100 µL per 10g of body weight. Mice were injected with tamoxifen 4 consecutive days in a row >1 week prior to analysis, and subsequent the day prior to injury injected with tamoxifen again to ensure recombination of the floxed allele (to total 5 injections).

### Muscle injury

Skeletal muscle injury was induced via tibialis anterior intramuscular of 50 µL 10µM cardiotoxin or 50 µL 1.2% weight/volume of barium chloride (BaCl_2_) using 27-gauge needle. Mice were first anesthetized using a ketamine/xylazine cocktail and intramuscular injection of the respective toxin was induced using a 27-gauge needle. Muscle samples were collected at the respective timepoints after injury and analyzed as described below. The contralateral muscle always served as a non-injured control.

### Cross-sectioning for muscle analysis

Cardiotoxin or BaCl_2_ injured muscle and the respective contralateral non-injured controls were collected at the respective timepoints after injury, weighed, embedded in O.C.T. (Fisher, 4585) and flash-frozen to preserve muscle tissue. Samples were cross-sectioned at 10µm thickness using a Leica CM1850 cryostat set at −20°C. Muscle sections were placed on a Tek-slide (IMEB Inc., Cat#Y-9253) and processed for hematoxylin and eosin staining, immunofluorescence, or stored at −80 °C.

### Hematoxylin & Eosin staining

Muscle cross-sections were first placed in hematoxylin for 15 minutes, rinsed with gently running water for 1-2 mins, placed in eosin for 1 minute, placed in ethanol to dehydrate samples (70%, 95%, 100%, 1 minute each), then xylene for 2 minutes and covered using Permount and a glass cover slip.

### Immunofluorescence staining

For slides, samples were surrounded by tissue blocker pen (for muscle fibers and myoblasts, samples were processed in 24 or 48-well plates) and fixed with 4% paraformaldehyde (PFA) for 10 minutes at room temperature. Samples were washed 3 times with 1X PBS (pH 7.5, 5 mins per wash), incubated with 1X glycine (0.375g/50mL dissolved in 1X PBS) for 10 minutes, and washed again 3 times with 1X PBS. Samples were blocked for 1 hour at room temperature in blocking buffer (5% goat serum, 2% bovine serum album, 0.1% triton X-100, 0.1% sodium azide prepared in 1X PBS). Primary antibodies were diluted in blocking buffer and samples were stained with the primary antibody over-night at 4 °C. Samples were calibrated to room temperature, washed with 1X PBST (1% tween-20 in 1X PBS) 3 times at room temperature (5-minute incubations). Samples were incubated with secondary antibodies and DAPI for 1 hour at room temperature, washed 3X with 1X PBST, and a drop of Permount was added and a coverslip placed on top to preserve fluorescence.

#### Primary antibodies

| Name | Catalog # | Host |
| --- | --- | --- |
| Anti-Pax7 | DSHB Cat# RRID:AB_2299243 | msIgG1 |
| Anti-Dystrophin | Abcam Cat# ab15277, RRID:AB_301813 | Rabbit |
| Anti-Myogenin | DSHB Cat# F5D, RRID:AB_2146602 | msIgG1 |
| Anti-Type IIA | DSHB Cat# SC-71 | mIgG1 |
| Anti-Type IIB | DSHB Cat# BF-F3 | msIgM |
| Anti-Type I | DSHB Cat# BA-D5 | msIgG2b |
| Anti-MF20 (MyHC) | DSHB Cat# MF 20, RRID:AB_2147781 |  |
| Anti-Ki67 | Abcam Cat# ab15580, RRID:AB_443209 | Rabbit |
| Anti-Myod | Santa Cruz Biotechnology Cat# sc-377460, RRID:AB_2813894 | msIgG2b |
| Anti-eMYHC | DSHB Cat# F1.652, RRID:AB_528358 | msIgG1 |
| Anti-Laminin | Sigma Cat# L9393, RRID:AB_477163 | Rabbit |

#### Secondary antibodies

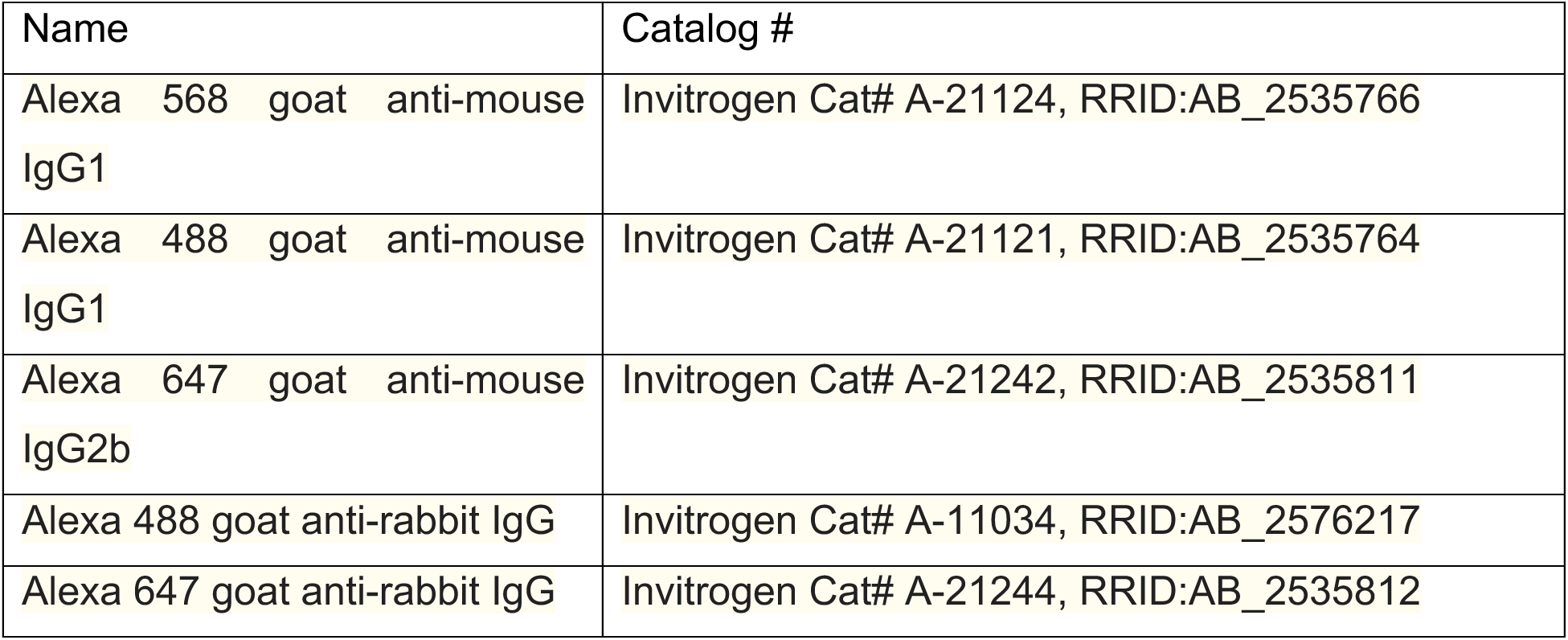

| Name | Catalog # |
| --- | --- |
| Alexa 568 goat anti-mouse IgG1 | Invitrogen Cat# A-21124, RRID:AB_2535766 |
| Alexa 488 goat anti-mouse IgG1 | Invitrogen Cat# A-21121, RRID:AB_2535764 |
| Alexa 647 goat anti-mouse IgG2b | Invitrogen Cat# A-21242, RRID:AB_2535811 |
| Alexa 488 goat anti-rabbit IgG | Invitrogen Cat# A-11034, RRID:AB_2576217 |
| Alexa 647 goat anti-rabbit IgG | Invitrogen Cat# A-21244, RRID:AB_2535812 |

### Imaging

All H&E and immunofluorescence samples were captured using a 20X or 10X objective on the Leica DM 6000B microscope. For each timepoint and pair, at least 4 images/genotypes were collected for analysis. Entire muscle images used for MuscleJ fiber type analysis were captured using the 10X objective on the EchoRevolution.

### RNA isolation and qRT-PCR

RNA for *in vitro* culture experiments was isolated at the indicated timepoints using TRIzol per the manufacturer’s instructions. RNA was resuspended in nuclease-free dH_2_O and measured using a Nano-drop. For reverse transcription reactions, 1µg of RNA/sample was diluted to a total volume of 10 µL/sample, and generation of complementary DNA was mediated by M-MLV reverse transcriptase following the manufacturers’ instructions (ThermoFisher, Cat#28025021). Final samples were diluted to a volume of 200µL prior to RT-PCR analysis. qRT-PCR was performed using FastStart Essential DNA Green Master Mix (Roche, Cat#06924204001) on a Roche Light cycler 96 in 96 well plates. Relative expression was measured using the 2^-DDCt^ method, and samples were normalized to *actin* expression.

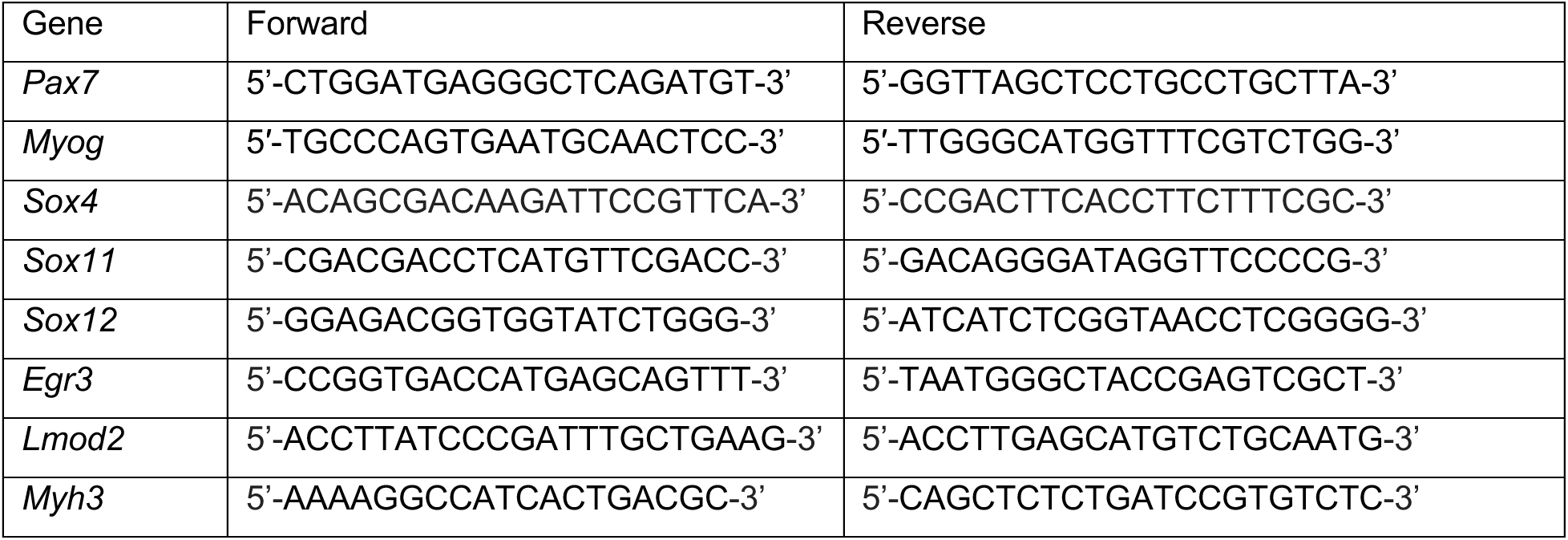

### Muscle fiber isolation and culture

For *ex vivo* muscle fiber culture, extensor digitorum longus (EDL) was carefully dissected from each mouse and digested with 2mg/mL of type I collagen in DMEM for 1 hour at 37°C with gentle inversions every 5 minutes. Individual muscles were transferred to pre-warmed DMEM to stop the digestion and triturated using a glass pipette to dissociate individual muscle fibers. Fibers were fixed immediately after digestion and dissociation (0 hour) or cultured in 20% FBS, 4 ng/mL bFGF, 1% penicillin-streptomycin in DMEM (Gibco) for 72 hours at 37°C with 5% CO_2_. For *in vitro* analysis of *Pax7*^CreER^ muscle fibers, fibers were cultured for 72 hours in medium containing 4-hydroxytamoxifen (4-OH, 0.4µM, Sigma) or methanol (MeOH, 1:1000). Muscle fibers were then fixed and processed as described above in 48-well plate.

### Primary myoblast isolation and culture

Primary myoblasts from *Pax7*^CreER^ or *Rosa26*^CreER^ litters were isolated using a modified protocol as described(57). Briefly, hindlimb muscles were dissected, placed in cold 1X PBS, washed 3X with 1X PBS to ensure all hair was removed, and gently cut by scraping in the direction of the muscle fiber to break up the tissue. Connective tissue and fat were removed, dissected samples were placed in 10 mL digestion buffer per mouse (700 U/mL type II collagenase, 10% horse serum, 1% penicillin-streptomycin in Ham’s F-10) and incubated at 37°C, 100rpm for 1 hour. Samples were neutralized with wash buffer (10% horse serum, 1% penicillin-streptomycin in Ham’s F-10), centrifuged at 500xg for 5 minutes, aspirated to leave 10mL medium, and incubated with 100 U/mL type II collagenase and 1U/mL dispase II at 37°C, 100rpm for 30 minutes. Samples were run through an 18-gauge needle 10X to break up additional pieces of muscle, run through a 40µm filter, centrifuged, supernatant aspirated and resuspended in 2 mL myoblast growth medium (20% FBS, 4 ng/mL bFGF, 1% penicillin-streptomycin in Ham’s F-10) and added to 10cm dish/sample containing myoblast growth medium. Cells were cultured for 48 hours to allow for myoblast proliferation, collected and digested with 0.025% trypsin-EDTA (Gibco), and pre-plated in 10cm dish for 45 minutes to allow for fibroblasts to attached. Supernatant from each plate was collected and placed on rat-tail collagen-coated plastic 10cm dishes, cultured for 24-48 hours, then collected and allowed to expand on non-coated plates to generate highly pure myoblasts. For proliferation, myoblasts were seeded on 24-well Matrigel coated plates in equal concentrations and evaluated after 24 hours. To induce differentiation, myoblasts were seeded in equal concentrations in 24-well Matrigel coated plates, grown for 24 hours in proliferation medium, switched to differentiation medium (2% horse serum, 1% penicillin-streptomycin in DMEM), and collected or analyzed at the indicated timepoints. For mixed cell cultures, cells collected immediately after isolation were plated on Matrigel coated plates in equal concentrations and grown in myoblast growth medium for 24 hours and then induced to differentiate via serum starvation (2% horse serum, 1% penicillin-streptomycin in DMEM) with either 4OH or MeOH. Samples +4OH or +MeOH were compared for both control mice or the experimental. However, samples pictured and analyzed *Pax7*^CreER^;Sox11^floxed/floxed^ or *Rosa26*^CreER^;Sox11^floxed/floxed^ cultured with MeOH or 4OH. Samples were then collected for RNA isolation or immunofluorescence as described.

### Muscle fiber cross sectional area and image analysis

For muscle fiber cross-sectional area analysis (CSA), dystrophin or laminin was used to delineate fiber boundary (as indicated). For all quantifications, images taken on 10X objective were dragged into Fiji and analyzed using MyoSight (67). For injured muscle, only muscle fibers with at least 1 centrally located nucleus were counted. For each pair of mice, >3 images were analyzed spanning the muscle section to yield >300 fibers/mouse analyzed. The frequency distribution is the average of at least 3 pairs of mice. For fiber type and CSA analysis, images of the entire muscle section were taken, loaded into MuscleJ and run to detected Type IIA and Type IIB (thus, Type IIX is inferred). Pax7 and Myogenin-positive cells per field of view is quantified as the average of the number of Pax7 or Myogenin-positive cells over 3-4 images/mouse for minimum of 3 biological replicates.

### Quantification and statistical analysis

Samples were plotted using GraphPad Prism 8. For all samples, at least 3 pairs were used. All samples are litter, age and gender matched. Unpaired t-test in Prism was used to analyze differences between genotypes.

## Results

### Identification of *Sox11* expression in myogenic progenitors

To identify key regulatory factors, pathways and mechanisms that stabilize cell identity and cell fate transitions, we first probed our previously published scRNA-seq dataset to determine genes that were uniquely enriched in the subsets of MuSCs (60). We found that *Sox11* expression is restricted to the MuSC population in cells profiled from non-injured and injured muscles at various timepoints throughout the regenerative process (Fig. S1A). We confirmed the expression on an aggregated scRNA-seq dataset (68), and found that indeed *Sox11* expression is mostly detected in the MuSC population (Fig. 1B, C). Additional analysis of published datasets (41, 50, 69) further showed that *Sox11* expression is specifically enriched in differentiating MuSCs (Fig. 1D) and this was corroborated by pseudo-temporal analysis (Fig. S1B). Quantitative qPCR analysis confirmed an increased expression of Sox11 during primary myoblast differentiation, along with Myod and Myog (Fig. 1E). This specificity was unique to *Sox11*, as *Sox4* expression gradually decline during differentiation (Fig. 1E).

We also employed *in silico* scTenifoldKnk to predict perturbations in gene regulatory networks that would be caused by knock-out of *Sox11* (65). Consistent with the expression pattern of Sox11 during MuSC differentiation, the top predicted gene targets impacted by knock-out of Sox11 are *Myh3, Lmod2, Mylk4, Myh8,* and *Mymk*, among others (Table S1). Many of these genes are developmentally expressed myosins and known regulators of newly regenerated muscle fibers (70). Together, these preliminary analyses identified *Sox11* as a novel marker and potential regulator of MuSC differentiation.

### *Sox11* expression is reduced with age in MuSCs, associated with changes in chromatin conformation and accessibility

Given that MuSC function declines with age (71, 72, 73), we sought to determine the cell autonomous and non-cell autonomous factors that are linked to age-related functional decline. To this end, we employed the 10X Chromium platform to garner the dynamics of transcriptional profiles of all mononuclear cells from regenerating hindlimb muscles of young (< 4 months) and old (> 22 months) male mice at 7 days post injury (DPI) (Fig. 2A). We processed both samples together, performed dimensionality reduction, clustering and UMAP embedding of the two samples (Fig. 2B). This yielded 7,874 gene expression profiles from young muscle samples and 6,013 gene expression profiles from old muscle samples. We identified known muscle cell populations based on marker gene expression and labeled them accordingly. This included a fibro-adipogenic progenitor (FAP) population expressing *Pdgfra* and *Postn*, 3 macrophage populations expressing *Cd68* and *Mrc1*, endothelial cells (ECs) enriched for *Pecam1*, smooth muscle cells (SMCs) expressing *Rgs5,* and a muscle stem cell (MuSC) population enriched for *Pax7* (Fig. 2C, D). These clusters recapitulated known cell populations involved in regeneration and allowed us to further probe any subtle differences in cell dynamics and gene expression that cannot be determined at the global population level.

**Figure 2.**
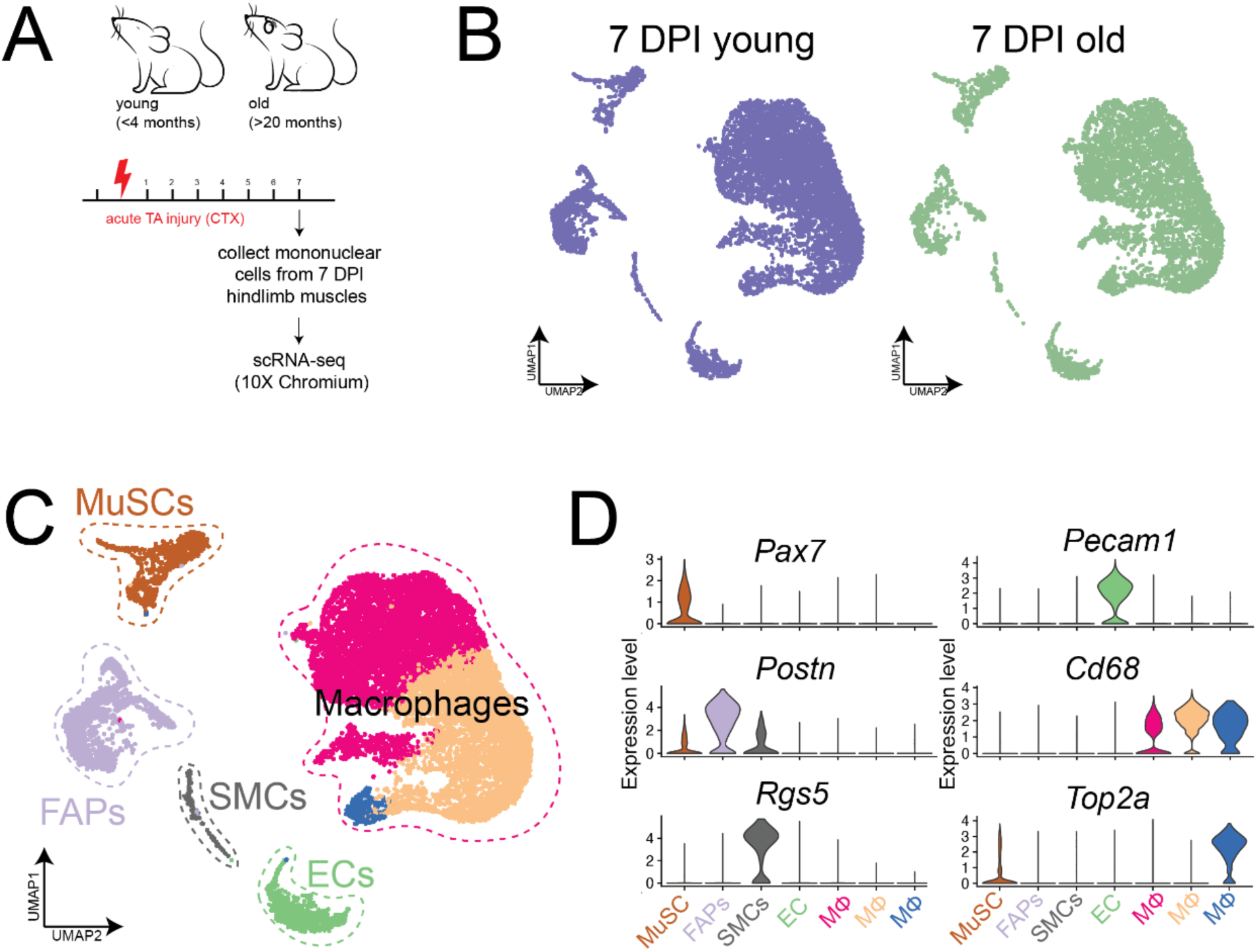
Single-cell RNA-sequencing of young and old mononuclear cells from 7 DPI injured muscle. **A.** Experimental outline: Mononuclear cells from hindlimb muscle from young (<4-month-old, n=3) and old (>20-month-old, n=3) male mice injured with CTX, collected at 7 DPI, and processed for scRNA-seq using the 10X Chromium Platform. **B.** UMAP of cells clustered together and separated based on young (left panel), and old (right panel) samples. **C.** UMAP and clustering results of cells combined from both young and old mice. Colored based on clusters identified. **D.** Violin plots for selected top marker genes used to label each cluster.

For the purpose of this study, we focused on the MuSC population and recluster the cells to determine any subtle changes (Fig. 3A, B). We subsequently evaluated gene expression differences based on age in the MuSC population and found that *Pax7* was relatively similar between the two ages, while markers associated with differentiation (*Myod1* and *Myog*) were significantly reduced in the MuSC subset. Interestingly, *Sox11* expression was also significantly reduced. Thus, *Sox11* expression, which increases as MuSCs differentiate, may also be reduced as a consequence of the delayed kinetics associated with age-related functional decline (71).

**Figure 3.**
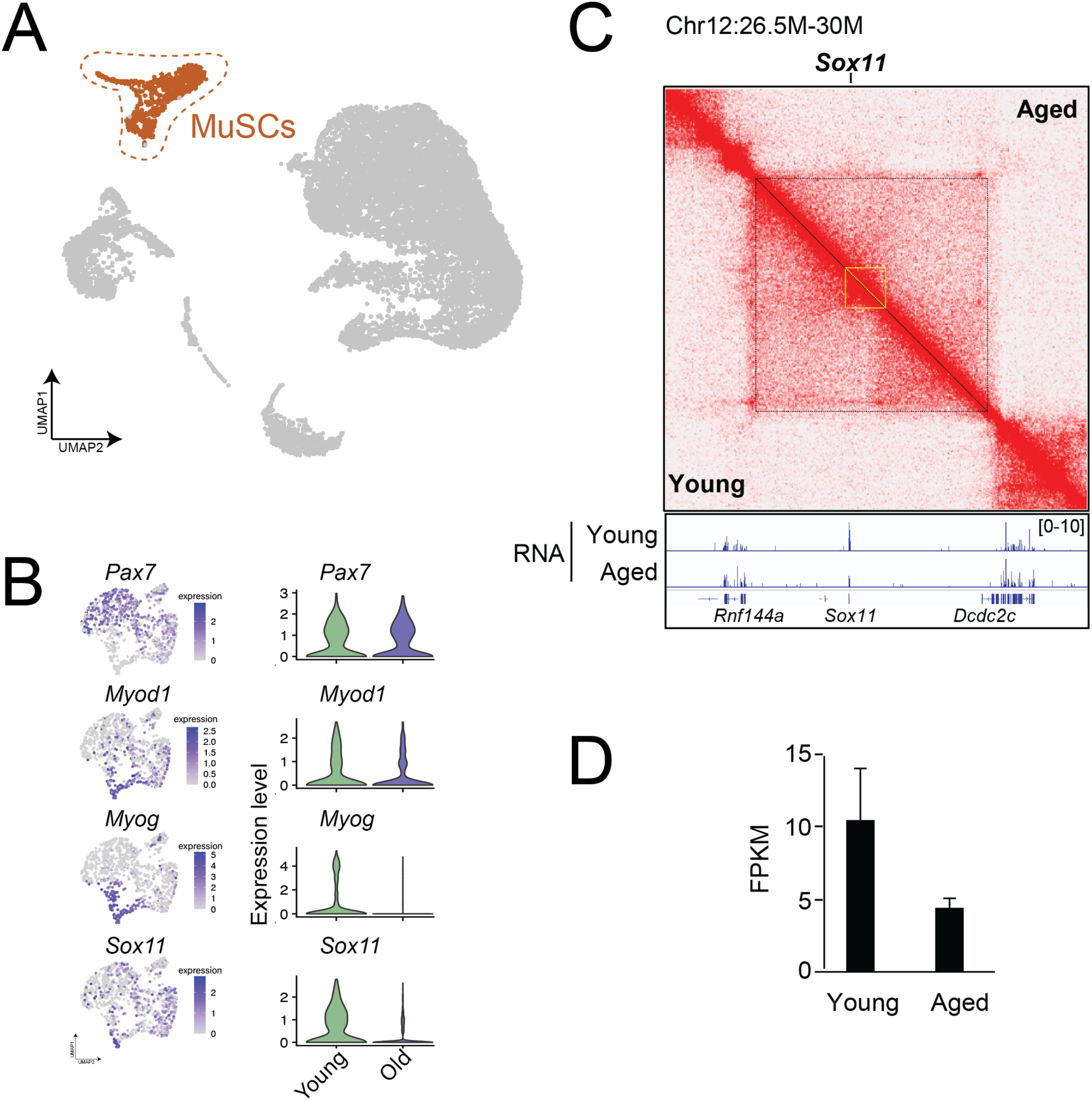
Reduced *Sox11* expression and 3D-genome alterations at *Sox11* locus in muscle stem cells with age. **A.** UMAP of all cells from young and old mice, MuSCs highlighted in color. **B.** Sub-setting on MuSCs and gene expression plots (left panel) and violin plots (based on age) identifies various differentiation-related transcripts are reduced with age identifies significantly reduced *Sox11* expression in MuSCs from old mice. **C.**Top: comparison of Hi-C contact maps (10-kb resolution) surrounding Sox11 locus between young and aged FISCs. The yellow triangle indicates the sub-TAD harboring Sox11 locus. Bottom, genome browser tracks showing the RNA-seq in young and aged FISCs. **D**. Bar graph showing the mean FPKM values of Sox11 in young and aged FISCs. n = 3 for each group.

Recent findings indicate that the 3D structure of the genome reorganizes and underpins the transcriptome remodeling during SC aging (74, 75, 76). Thus, we leveraged our recently published Hi-C datasets (74) to investigate the 3D organization around the *Sox11* locus. In freshly isolated MuSCs (FISCs) from young mice, a prominent topologically associated domain (TAD) structure harboring *Sox11* locus was observed, and it was largely maintained in aged FISCs (Fig. 3C). *Sox11* was found in the boundary region of one sub-TAD which was delineated by an upstream CTCF binding site (Fig. 3C). Notably, the abundance of the *Sox11* containing sub-TAD decreased in aged FISCs (Fig. 3C), which is in agreement with our observations of reduced *Sox11* expression in aged samples. Consistently, *Sox11* expression was substantially decreased in the bulk RNA-seq data (Fig. 3D), which is in agreement with our observations in scRNA-seq analysis. The above data indicate that the decline in Sox11 expression during MuSC aging is associated with 3-dimensional genome reorganization at the *Sox11* locus.

### *Sox11* is dispensable for adult muscle stem cell function and muscle regeneration in vivo

To determine if the dynamic expression of *Sox11* during myogenesis and aging is underlies a function of Sox11 protein in adult MuSC function, we generated a *Pax7*^CreER^;*Sox11*^fl/fl^ inducible mouse model to specifically delete *Sox11* in Pax7+ MuSCs and their progeny upon the administration of tamoxifen (Fig. 4A). To evaluate MuSC function upon loss of *Sox11*, we used a model of acute injury to evaluate MuSCs’ ability to activate, proliferate, and differentiate *in vivo* (77). To assess the impact that loss of *Sox11* has on the ability of the MuSC pool to efficiency repair damaged fibers, we administered tamoxifen to control (*Pax7*^CreER^;*Sox11*^fl/+^, *Sox11*^fl/fl^ or *Sox11*^fl/fl^) and *Pax7*^CreER^;*Sox11*^fl/fl^ (Sox11-pKO), induced acute injury to the tibialis anterior (TA) muscles, and analyzed muscle regeneration at discrete timepoints throughout the regenerative process (Fig. 4A).

**Figure 4.**
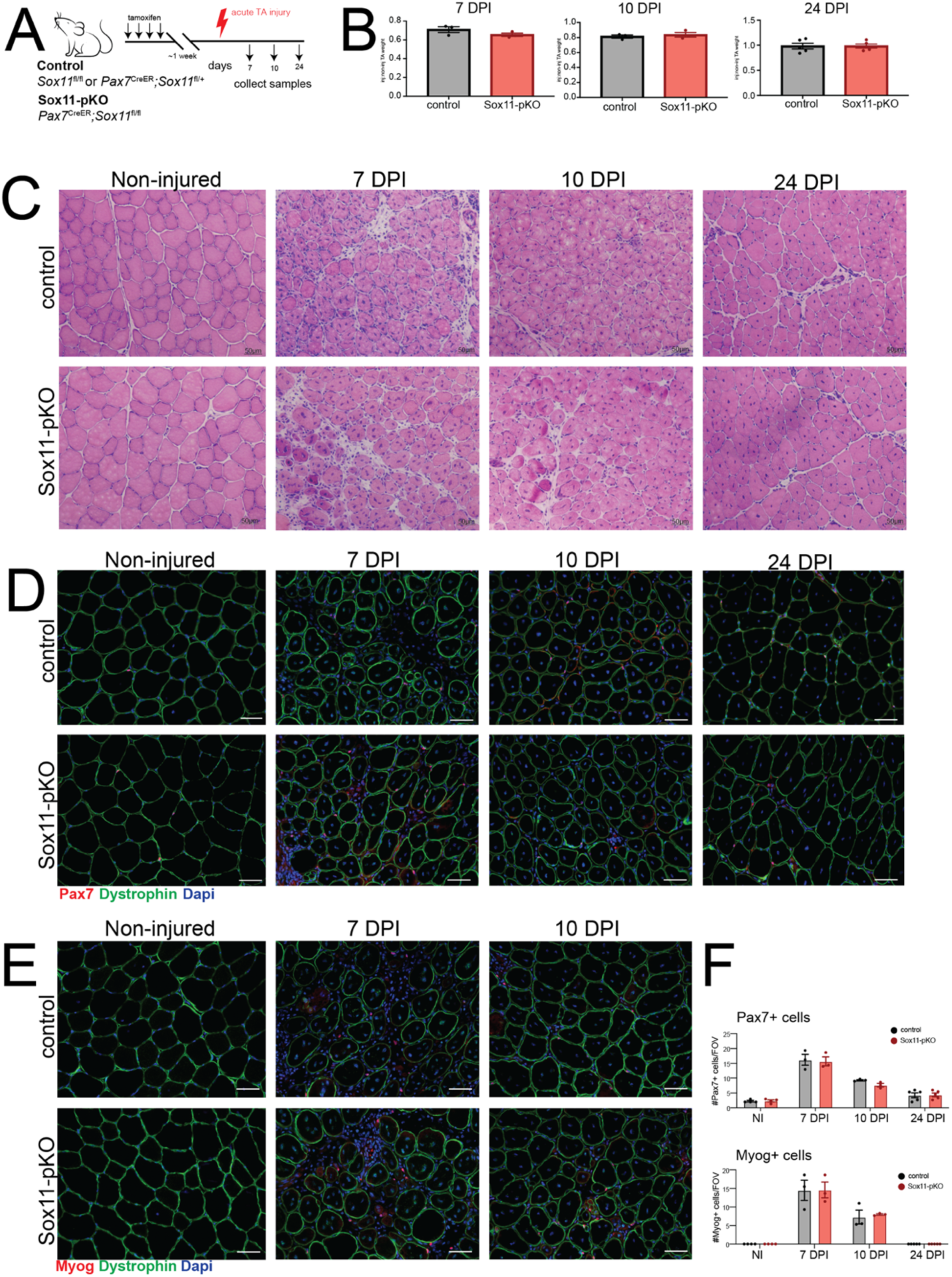
Analysis of muscle fiber area and type for Sox11-pKO mice. **A.** Frequency distribution pots for CSA of TA muscle fibers from non-injured, 7, 10, and 24 DPI mice from control and Sox11-pKO mice. Measurements binned in 400 um^2^ bins. **B.** Representative immunofluorescence images to detect fiber Type IIA, Type IIB, Dystrophin and nuclei (DAPI) on non-injured muscles from control and Sox11-pKO mice (top panel), output from MuscleJ (bottom panel). **C.** Frequency distribution pots for CSA of TA muscle fibers from non-injured muscle, separated by inferred fiber type as analyzed from MuscleJ. Measurements binned in 400 um^2^ bins. **D.** Representative immunofluorescence images to detect fiber Type IIA, Type IIB, Dystrophin and nuclei (DAPI) on 24 DPI muscles from control and Sox11-pKO mice (top panel), output from MuscleJ (bottom panel). **E.** Frequency distribution pots for CSA of TA muscle fibers from 24 DPI muscle, separated by inferred fiber type as analyzed from MuscleJ. Measurements binned in 400 um^2^ bins.

No significant differences in muscle mass recovery (weight) were observed between the two genotypes at 7, 10, and 24 DPI, suggesting that Sox11-pKO MuSCs are functionally competent to regenerate injured muscles (Fig. 4B). Consistently, normal muscle morphology was observed in control and Sox11-pKO mice, with no clear differences in regenerated myofibers between the two genotypes at 7 and 10 DPI (Fig. 4C). By 24 DPI, both control and Sox11-pKO injured muscle appeared nearly fully repaired (Fig. 4C). Additional analysis of muscle fiber cross-sectional area (CSA) at 7, 10 and 24 DPI indicated that control and Sox11-pKO regenerated fiber area were comparable at each timepoint (Fig. S2A). While at 10 DPI, Sox11-pKO had significantly more fibers of 1200-2800 um^2^, this modest change in size distribution was resolved by 24 DPI (Fig. S2A).

To determine if loss of *Sox11* expression alters the activation, proliferation, or differentiation status of MuSCs, we evaluated both Pax7 and Myogenin expression via immunofluorescence on muscle sections from each injury timepoint (Fig. 4D, E). Both control and Sox11-pKO had comparable numbers of Pax7+ cells and Myog+ cells per field of view (FOV) in non-injured and at 7, 10 and 24 DPI muscle (Fig. 4F). Specifically, both genotypes had nearly 20 Pax7+ cells/FOV at 7 DPI, which decreased to 10 and 5 by 10 and 24 DPI, respectively. The number of Myog+ cells/FOV reached an average of 15 at 7 DPI for both genotypes and subsequently decreased to less than 10 by 10 DPI.

Since another Sox family member, Sox6, regulates slow-muscle fiber genes, we evaluated fiber type distribution of non-injured and regenerated muscle at 24 DPI (53, 54). However, fiber type and size distribution at 24 DPI were similar between control and Sox11-pKO, indicating that *Sox11* is required for muscle fiber type determination after acute TA muscle injury (Fig. S2B). Thus, Sox11 appears dispensable for MuSCs function in response to acute injury.

Multiple rounds of injury can exacerbate any subtle differences in regenerative capacity while also assessing the ability of MuSCs to self-renew (78). To determine if *Sox11* knockout changes the proportion of self-renewing and differentiating MuSCs, we performed repetitive rounds of injury and analyzed muscle at 7 days after the 3^rd^ injury (Fig. S3A). Sox11-pKO recovery TA weight was significantly greater than control, although gross muscle morphology and regeneration appeared comparable between control and Sox11-pKO (Fig. S3B, S3C). Interestingly, we observed significantly more Pax7+ cells per FOV in Sox11-pKO at 7DPI after the 3^rd^ injury, which was greater than 10/FOV in Sox11-pKO compared to an average of 7.5 Pax7+ cells/FOV in control (Fig. S3C). We observed a marginal difference that did not research statistical difference in the number of Myog+ cells/FOV between the two genotypes (Fig. S3C). Thus, loss of *Sox11* may slightly delay the differentiation of MuSCs which is exacerbated by multiple rounds of injury. Nonetheless, Sox11 is not absolutely required for adult MuSC function in response to acute injury or for efficient regeneration after multiple rounds of injury.

### *Sox11* is expendable for muscle stem cell function *in vitro*

The observation that Sox11-pKO mice had a greater proportion of Pax7+ cells per FOV after multiple rounds of injury (Fig. S3) prompted us to further probe if Sox11 KO influences MuSC function cell autonomously. We isolated muscle fibers and their associated MuSCs from Sox11-pKO mice and cultured them in the presence of 4-hydroxy tamoxifen (4-OH) to induce recombination of the floxed allele or MeOH (vehicle control). We fixed fibers at 0 hours and 72 hours in culture and used immunofluorescence to detect Pax7 and Myod (Fig. 5A). However, there was no significant differences between the MeOH or 4OH treated muscle fibers, with both genotypes averaging approximately 8 cells/cluster (Fig. 5B).

**Figure 5.**
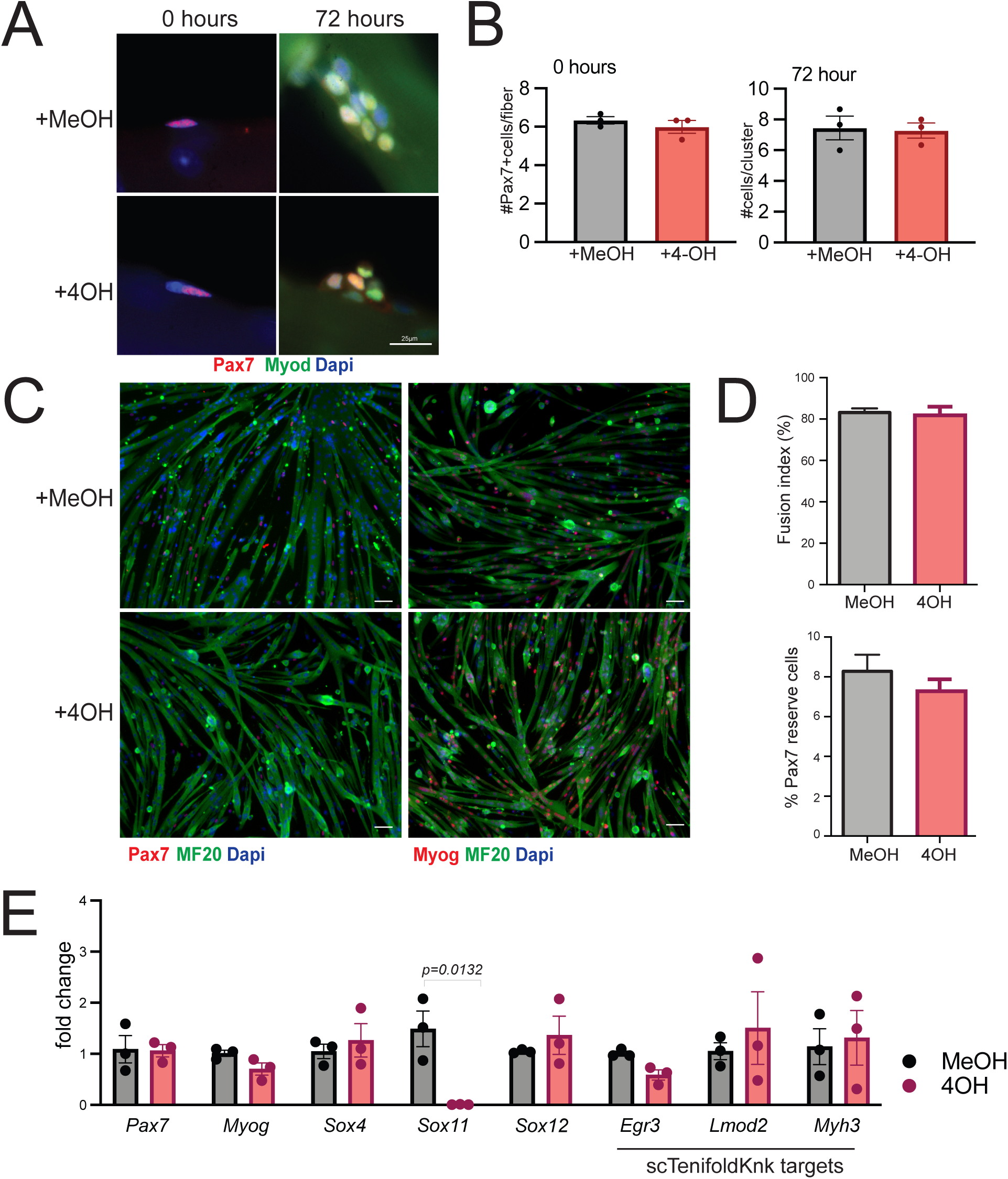
*In vitro* analysis of myogenic function Sox11-null myoblasts. **A.** Representative immunofluorescence images to detect Pax7, Myod1 and nuclei (counterstained with dapi) of myofibers and their associated MuSCs isolated from *Pax7*^CreER^;*Sox11*^fl/fl^ EDL muscle and fixed at 0 hours (left panel) or cultured for 72 hours with vehicle (MeOH) or 4-OH (right panel). **B.** Quantification of the number of Pax7+ cells/myofiber at 0 hours (left graph) and the number of cells/clusters at 72 hours in culture (right graph); related to panel A. **C.** Primary myoblasts isolated from *Pax7*^CreER^; *Sox11*^fl/fl^ were cultured with MeOH (top panel) or 4-OH (bottom panel) plated at equal concentrations and induced to differentiate via serum starvation for 4 days to evaluate myogenic potential. Representative immunofluorescence images to detect Pax7, MF20, nuclei (DAPI) shown in the left panel and Myog, MF20, and nuclei (DAPI) are shown in the right panel. **D.** Quantification of the fusion indices of MeOH and 4OH treated myoblasts, represented as the percent of nuclei fused into myotubes (top graph) and Pax7 ‘reserve’ cells, counted as the total number of Pax7+ cells per field of view for each biological replicate (bottom graph). **E.** Relative expression as measured by qRT-PCR for MuSC marker *Pax7* and differentiation marker *Myog*, members of the SoxC sub-family (*Sox4, Sox11* and *Sox12*), and genes predicted to be reduced by scTenifoldKnk (*Egr3*, *Lmod2*, *Myh3*). Scale bars: 25 µm in A, 50 µm in C.

To further evaluate MuSC proliferation and differentiation, we isolated primary myoblasts from Sox11-pKO mice and cultured them in the presence of 4OH or MeOH. We confirmed recombination of the floxed allele (Fig. S4A), and subsequently evaluated proliferation and differentiation. Sox11-pKO myoblasts treated with MeOH or 4-OH had comparable numbers of Pax7+/Ki67+ cells and comparable numbers of Myog+ cells (Fig. S4B) suggesting no overt proliferative and differentiation defects. Given that *Sox11* was enriched during differentiation, we induced differentiation through serum starvation of Sox11-pKO myoblasts treated with 4OH or MeOH (vehicle control). Immunofluorescence on differentiated cells to detect Pax7 or Myog and MF20 suggested control (+MeOH) and Sox11-null (+4-OH) myoblasts differentiated similarly (Fig. 5C). Control and Sox11-null myoblast fusion indices both reached 80% and the number of ‘reserve’ Pax7+ cells, which may represent self-renewal capacity of myoblasts (79), were comparable with MeOH treated reaching near 8% and 4OH treated reaching near 7% Pax7 reserve cells (Fig. 5D). Gene expression analysis on RNA isolated from differentiated myoblasts confirmed that *Sox11* expression was ablated while *Pax7,* and *Myog* were unperturbed in the knockout cells (Fig. 5E). The expression of the other SoxC genes, *Sox4* and *Sox12*, were unchanged between the two conditions, excluding the possibility that a compensatory increase in expression of other Sox members may have accounted for the lack of phenotype (Fig. 5E). We also evaluated the expression of scTenifoldKnk Sox11-KO genes that were predicted to be dysregulated upon Sox11 knock-out and found no significant differences in their expression upon loss of *Sox11* (Fig. 5E). In conclusion, *Sox11* is not required for MuSC self-renewal or differentiation *in vitro*.

### Loss of *Sox11* in muscle progenitors minimally impacts embryonic muscle development

Although we determined that *Sox11* is unnecessary for adult MuSC function, many of the Sox family of transcription factors are highly expressed during development and are required for proliferation and survival as well as differentiation of various embryonic cell types (35, 36). We therefore evaluated the requirement of *Sox11* for muscle progenitor function during development by generating *Myod1*^Cre^; *Sox11*^fl/fl^ mice to specifically delete Sox11 in Myod1+ muscle progenitors and their progeny. Control (*Myod1*^Cre^;*Sox11*^f*l*/+^, *Sox11*^f*l*/fl^ or *Sox11*^f*l*/+^) and *Myod1*^Cre^;*Sox11*^f*l*/fl^ (Sox11-mKO) mice were born in Mendelian ratios and had comparable body weights (Fig. S5A). There were no observable differences in muscle morphology or CSA by 3 months of age between control and Sox11-mKO mice (Fig. S5B, S5C). Therefore, although Sox11 is important for tissue specification and organogenesis, it does not appear overtly necessary for normal muscle development. To further probe the impact that loss of *Sox11* in muscle progenitors has on MuSC function, we induced acute injury via intramuscular injection of CTX and analyzed samples at 5.5 DPI (Fig. S5A). Control and Sox11-mKO had similar TA recovery weights and no obvious differences in muscle morphology and regeneration 5 DPI (Fig. S5B, S5C). Immunofluorescence on muscle sections to detect Pax7 or Myog and Laminin supported the lack of observable differences between control and Sox11-mKO, with control and Sox11-mKO having an average of 35 Pax7+ cells/FOV and an average of approximately 20 Myog+ cells/FOV for both genotypes (Fig. S5C). Thus, *Sox11* is dispensable for the establishment and function of the MuSC pool.

### Loss of Sox11 does not accelerate age-related functional decline

Since we identified *Sox11* expression to be reduced in old MuSCs from 7 DPI and found age-related changes in chromatin conformation (Fig. 3), we sought to evaluate whether loss of *Sox11* exacerbates any age-related regenerative function (34, 80). We therefore injured the TA muscles of old (∼22 month old) control and Sox11-mKO mice and analyzed samples at 7 DPI (Fig. 6D). Both control and Sox11-mKO mice regenerated efficiently, as evidenced by immunofluorescence to detect Pax7 or Myog and Laminin which were indistinguishable (Fig. 6E). Quantification of the number Pax7+ or Myog+ cells per FOV indicated age did not lead to any obvious alterations in MuSC’s ability to repair damaged fibers, both having an average of approximately 35 Pax7+ cells/FOV and 15 Myog+ cells/FOV at 7 DPI (Fig. 6F). While measurement of muscle fiber CSA suggested that aged Sox11-mKO had significantly fewer fibers in the range of 2000-2400um^2^, regenerated fiber CSA was similar between control and Sox11-mKO (Fig. S5D). *Ex vivo* fiber and MuSC culture results were consistent with *in vivo* results, with no significant differences observed in Pax7+ cells per myofiber or cluster size (Fig. 6G, 6H). Therefore, although we identified reduced *Sox11* expression in MuSCs from aged mice, loss of *Sox11* does not appear to accelerate any age-related regenerative capacity of MuSCs.

**Figure 6.**
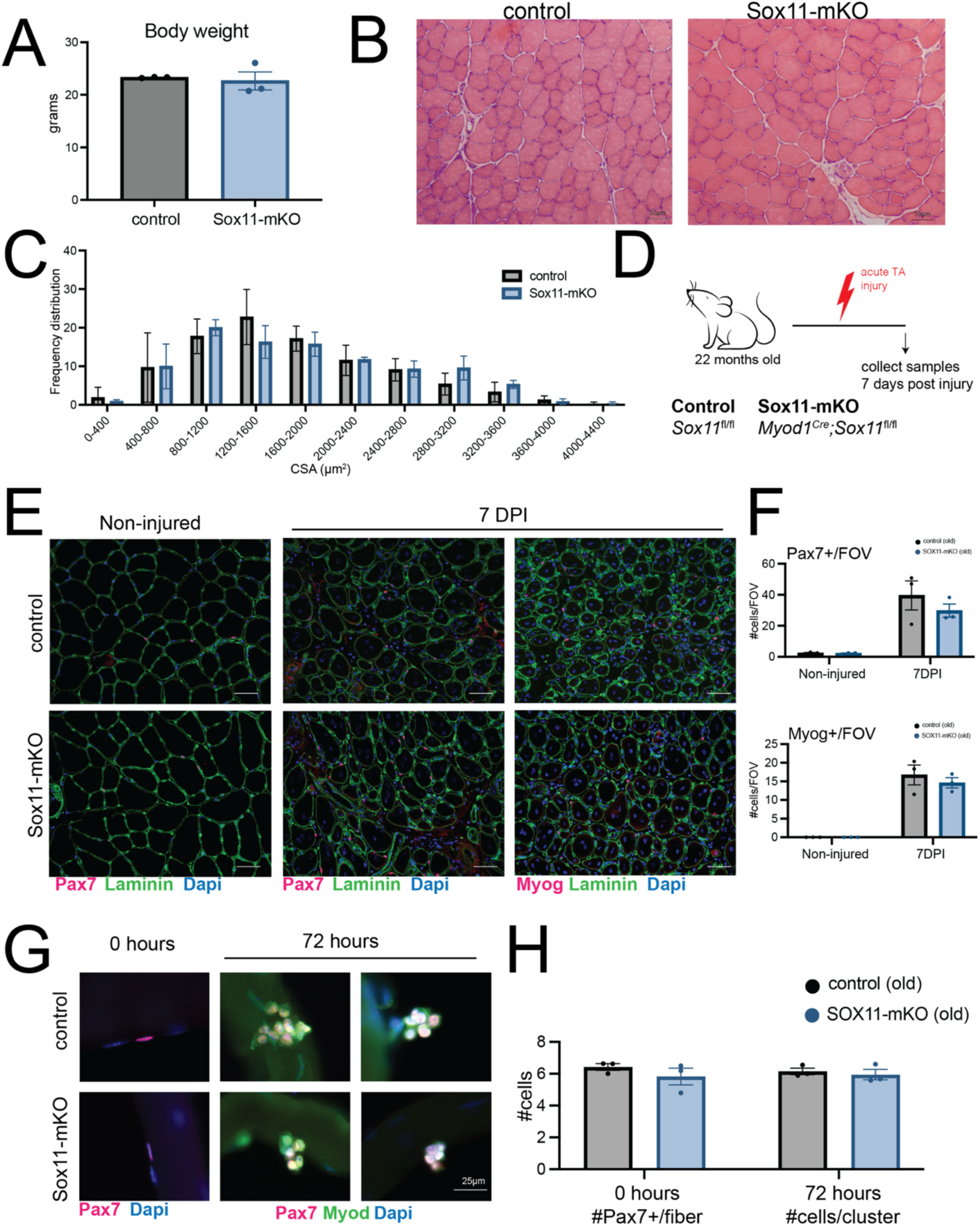
Requirement of Sox11 in muscle development and aging. **A.** Body weight of control and Sox11-mKO mice in grams. **B.** H&E staining and representative images of 10µm TA muscle cross-sections from control (top panel) and Sox11-mKO (bottom panel) non-injured samples, scale bars: 50 µm. **C.** TA muscle fiber CSA for non-injured control and Sox11-mKO mice, related to B. **D.** Experimental outline to evaluate the impact loss of Sox11 has on regeneration of aged (>20-month-old) mice. The TA muscle of control and Sox11-mKO mice were injured via intramuscular injection of CTX and collected at 7 DPI to evaluate regeneration**. E**. Representative images of immunofluorescence on TA muscle sections to detect Pax7 or Myog, Dystrophin, and nuclei (counterstained with DAPI) from control (top panel) and Sox11-mKO (bottom panel) old mice at 0 (non-injured), and 7 DPI, scale bars: 50 µm **F.** Quantification of the number of Pax7+ cells/FOV (top graph) and Myog+ cells/FOV (bottom graph) for control and Sox11-pKO mice. **G.** *Ex vivo* culture of muscle fibers isolated from EDL of old control (top panel) and Sox11-mKO (bottom) mice, fixed at 0 hours (left panel), or after 72 hours in culture (right panel) and stained to detect Pax7, Myod and nuclei (DAPI), scale bar 25 µm. **H.** Quantification of the number of Pax7+ cells/myofiber at 0 hours (left graph) and the number of cells/clusters at 72 hours in culture (right graph); related to panel G

### Sox11 is globally dispensable for muscle repair after acute injury

A recent study specifically performed scRNA-seq on nuclei isolated from muscle spindles and found *Sox11* expression to be unique to the sensory bag fibers (81). Furthermore, Sox11 is required for sensory neuron regeneration, and thus may more broadly play a role in regeneration (41, 50). To understand if Sox11 is globally required for adult muscle regeneration, we crossed *Rosa26*^CreER^ mice with *Sox11*^fl/fl^ mice to delete *Sox11* in all cell types upon the administration of tamoxifen. We administered tamoxifen to control (*Rosa26*^CreER^;*Sox11*^fl/+^, *Sox11*^fl/fl^ or *Sox11*^fl/fl^) and *Rosa26*^CreER^;*Sox11*^fl/fl^ mice (Sox11-KO) (Fig. S6A). We induced acute muscle injury with BaCl_2_ and analyzed regeneration at 5, 7, 10 and 24 DPI (Fig. S6A). Control and Sox11-KO recovery TA weights were similar, suggesting broadly efficient muscle regeneration between the two genotypes (Fig. S6B). Cross-sections of non-injured and injured TA muscles and H&E staining revealed similar morphological characteristics between control and Sox11-KO and each regeneration timepoint (Fig. S6C). Additional analysis of TA muscle sections using immunofluorescence for Dystrophin, Pax7 and Myogenin further supported the relatively similar regenerative capacity of control and Sox11-KO (Fig. S6D, S6E). Quantification of the number of Pax7+ and Myog+ cells per FOV indicated control and Sox11-KO mice were indistinguishable across the timepoints assayed (Fig. S6F). We therefore concluded that Sox11 is globally dispensable for adult muscle regeneration. Thus, while Sox11 is required for a variety of developmental processes, its role in muscle function and repair response to acute injury is limited.

## Discussion

The SOX family TFs play diverse roles in regulation of cell identity, self-renewal, and differentiation through their modulation of various transcriptional programs (82). For example, during skeletogenesis, Sox11 was found to stabilize nuclear β-catenin thereby promoting canonical WNT signaling to secure cell fate and shown to specifically regulate WNT-related pathway genes required for mesenchyme specification and organogenesis (83, 84). During muscle development, WNT ligands are secreted from the neural tube to promote myogenesis of the adjacent somite (31, 85). In resting adult muscle, the WNT signaling pathway is not active, but is activated in response to injury and *in vitro* during differentiation and increased WNT signaling impairs the regenerative potential of MuSCs (33, 34, 72, 86, 87). Interestingly, β-catenin-dependent WNT signaling opposes Notch signaling to promote MuSC differentiation and Sox11 was also shown to regulate the expression of Notch-related pathway genes (45, 47, 48, 88). We identified *Sox11* expression to be unique to differentiating MuSCs and reduced with age, consistent with age-dependent changes in chromatin conformation at the *Sox11* locus. We therefore hypothesized that Sox11 may play a role in the transcriptional regulation of MuSC function and fate decisions.

To this end, we employed a handful of genetic models to evaluate the requirement for Sox11 in muscle development and regeneration. To specifically evaluate the role for Sox11 in the adult muscle stem cell pool, we used a tamoxifen inducible MuSC specific model to delete *Sox11* in MuSCs. We sampled muscle at 7, 10 and 24 DPI to garner a complete picture of the regenerative process. The results indicate that control and Sox11-pKO exhibit comparable rates of regeneration. Thus, loss of *Sox11* does not impact the ability of MuSCs to activate, proliferate and differentiation in response to injury. These data were corroborated by *ex vivo* myofiber culture and *in vitro* differentiation of myoblasts. While Sox11-pKO muscles had an increased number of Pax7+ MuSCs at 7 days after a 3^rd^ muscle injury, muscle regeneration was relatively normal, with no other overt defects observed. Thus, Sox11 is not required for adult MuSCs function in response to one or multiple rounds of injury.

Since the Sox family of TFs are critically known for their development roles, we deleted Sox11 in muscle progenitors but found that muscle developed normally and maintained regenerative capacity into adulthood. As we identified age-related reduced *Sox11* expression and changes in chromatin conformation at the *Sox11* locus, we evaluated whether loss of *Sox11* impacts regenerative capacity with age. Surprisingly, we did not observe overt differences in regenerative capacity with age either. To outline the potential non-cell-autonomous requirement for Sox11 in adult skeletal muscle regeneration, we deleted Sox11 in all adult cells and induced acute muscle injury. However, assessment of the muscle regeneration after acute injury indicated that Sox11 is globally dispensable for muscle regeneration.

Given that a lack of Sox11 antibodies precluded protein analysis, we cannot exclude that the knock-out is not affected. It may also be that Sox4 and Sox11 play redundant roles in MuSC function, which will require future studies to clearly delineate. Although our data indicate that Sox11 is not required for normal MuSC activation, proliferation, and differentiation to repair muscle injury in response to acute injury, it remains to be determined if Sox11 plays a role under other conditions, such as nerve crush injury. Furthermore, the impact loss of Sox11 has on the aging MuSC phenotype will need to be explored, as loss of Sox11 does not exacerbate regenerative decline with age but may lead to changes in the MuSC aging phenotype. Nonetheless, our data provide the community with knowledge about the unique stage-specific expression and dispensable role of Sox11 in muscle development and acute muscle injury repair.

## Conclusions

We used scRNA-seq and 3D chromatin conformation assays to demonstrate unique enrichment of Sox11 expression in MuSCs and its stage/age-dependent expression dynamics, associated with changes in three-dimensional genome organization at the *Sox11* gene locus. Through a series of genetic assays using conditional knockout models, we found that Sox11 expression in myogenic and non-myogenic cells is dispensable for normal muscle development, MuSC regenerative function in response to injury in adulthood, and age-related muscle maintenance and regeneration in adulthood. Further studies to determine whether other SOX TFs can compensate for Sox11 may lend further insight into a potential regulatory role of Sox11 in myogenesis.

## Declarations

## List of abbreviations

4OH: 4-hydroxy-tamoxifen
BaCl_2_: Barium chloride
CSA: Cross-sectional area
CTCF: Transcriptional repressor CCCTC-binding factor
CTX: Cardiotoxin
DPI: Days post injury
ECs: Endothelial cells
FAPs: Fibro-adipogenic progenitors
FISCs: Freshly isolated muscle stem cells
FOV: Field of view
H&E: Hematoxylin & eosin
MeOH: Methanol
MRFs: Myogenic regulatory factors
MuSCs: Muscle stem cells
NICD: Notch intracellular domain
scRNA-seq: Single-cell RNA-sequencing
SMCs: Smooth muscle cells
Sox11-KO: *Rosa26*^CreER^;*Sox11*^fl/fl^ (+ tamoxifen *Rosa26*^CreER^;*Sox11*^-/-^)
Sox11-mKO: *Myod1*^Cre^;*Sox11*^fl/fl^ (in myogenic progenitors *Myod1*^Cre^;*Sox11*^-/-^)
Sox11-pKO: *Pax7*^CreER^;*Sox11*^fl/fl^ (+ tamoxifen in MuSCs *Pax7*^CreER^;*Sox11*^-/-^)
TA: Tibialis anterior
TAD: Topologically associating domain
TFs: Transcription factors
UMAP: Uniform manifold approximation

## Data availability

scRNA-seq data generated in this study can be found under the GEO accession number GSE226907. Previously published and used scRNA-seq data can be found under the accession numbers GSE150366 (scRNA-seq of MuSCs from non-injured, 5 and 10 DPI muscle) and GSE162172 (aggregated scRNA-seq data from skeletal muscle).

## Competing interests

The authors declare that they have no conflicts of interest with the contents of this article.

## Acknowledgments

We thank Dr. Veronique Lefebvre for kindly providing the Sox11-floxed mice, Dr. Timothy Ratliff and Dr. Gregory Cresswell for assistance with the 10X Genomics Platform, Dr. Phillip San Miguel and Purdue’s Genomics Core for RNA-sequencing, Dr. Luiz Brito for access to cluster computing, Dr. Jill Hutchcroft and Purdue’s Flow Cytometry and Cell Separation Facility for assistance with FACS data collection and analysis, and Jun Wu for technical support.

## Funding

This work was partially supported by grants from the National Institutes of Health to S. O. (F31AR077424) and S. K. (R01AR078695, R01AR079235, R01DK132819).

## Notes

### Competing Interest Statement

The authors have declared no competing interest.

