## Supplemental Material for "Sox11 is enriched in myogenic progenitors but dispensable for development and regeneration of skeletal muscle"

### Tables

**Table S1.** *scTenifoldKnk* prediction of top ranked genes dysregulated by Sox11 knokcout in muscle satellite cells*.*

| **Gene** | **distance** | **Z** | **FC** | **P.value** | **P.adj** |
| --- | --- | --- | --- | --- | --- |
| *Sox11* | 2.49E-06 | 9.40741082 | 8.47E+03 | 0.00E+00 | 0.00E+00 |
| *Myh3* | 1.77E-06 | 8.99923945 | 4.25E+03 | 0.00E+00 | 0.00E+00 |
| *Lmod2* | 6.42E-07 | 7.84827863 | 5.63E+02 | 2.23E-124 | 3.58E-121 |
| *Mylk4* | 2.59E-09 | 2.70212039 | 9.18E-03 | 9.24E-01 | 1.00E+00 |
| *Mybph* | 2.56E-09 | 2.69117503 | 8.93E-03 | 9.25E-01 | 1.00E+00 |
| *Casq2* | 2.47E-09 | 2.66406614 | 8.33E-03 | 9.27E-01 | 1.00E+00 |
| *Myh8* | 2.31E-09 | 2.61230442 | 7.29E-03 | 9.32E-01 | 1.00E+00 |
| *Mymk* | 2.31E-09 | 2.61035514 | 7.26E-03 | 9.32E-01 | 1.00E+00 |
| *Myom2* | 2.27E-09 | 2.59973918 | 7.06E-03 | 9.33E-01 | 1.00E+00 |
| *Tmem182* | 2.27E-09 | 2.59758595 | 7.02E-03 | 9.33E-01 | 1.00E+00 |
| *Kif4* | 2.18E-09 | 2.56527485 | 6.46E-03 | 9.36E-01 | 1.00E+00 |
| *Unc45b* | 2.16E-09 | 2.56050159 | 6.38E-03 | 9.36E-01 | 1.00E+00 |
| *Myo18b* | 2.13E-09 | 2.55061697 | 6.22E-03 | 9.37E-01 | 1.00E+00 |
| *Lmod3* | 2.12E-09 | 2.54457698 | 6.12E-03 | 9.38E-01 | 1.00E+00 |
| *Xirp2* | 2.08E-09 | 2.52991198 | 5.90E-03 | 9.39E-01 | 1.00E+00 |
| *Trdn* | 2.04E-09 | 2.51642624 | 5.69E-03 | 9.40E-01 | 1.00E+00 |
| *Klhl41* | 2.03E-09 | 2.51033576 | 5.60E-03 | 9.40E-01 | 1.00E+00 |
| *Hist1h2ap* | 1.89E-09 | 2.45846764 | 4.90E-03 | 9.44E-01 | 1.00E+00 |
| *Actc1* | 1.87E-09 | 2.44749006 | 4.76E-03 | 9.45E-01 | 1.00E+00 |
| *Kif11* | 1.85E-09 | 2.44171296 | 4.69E-03 | 9.45E-01 | 1.00E+00 |
| *Hist1h1b* | 1.85E-09 | 2.43979342 | 4.66E-03 | 9.46E-01 | 1.00E+00 |
| *Gm28653* | 1.81E-09 | 2.42554875 | 4.49E-03 | 9.47E-01 | 1.00E+00 |
| *Knl1* | 1.80E-09 | 2.41869322 | 4.41E-03 | 9.47E-01 | 1.00E+00 |
| *Egr3* | 1.78E-09 | 2.41221892 | 4.34E-03 | 9.47E-01 | 1.00E+00 |
| *Csrp3* | 1.78E-09 | 2.41054708 | 4.32E-03 | 9.48E-01 | 1.00E+00 |
| *Hmmr* | 1.66E-09 | 2.35641795 | 3.75E-03 | 9.51E-01 | 1.00E+00 |
| *Casz1* | 1.63E-09 | 2.34432181 | 3.63E-03 | 9.52E-01 | 1.00E+00 |
| *Myog* | 1.62E-09 | 2.34103138 | 3.60E-03 | 9.52E-01 | 1.00E+00 |
| *Kif15* | 1.62E-09 | 2.34004687 | 3.59E-03 | 9.52E-01 | 1.00E+00 |
| *Cenpf* | 1.58E-09 | 2.31918579 | 3.40E-03 | 9.54E-01 | 1.00E+00 |
| *Mymx* | 1.54E-09 | 2.30275057 | 3.26E-03 | 9.54E-01 | 1.00E+00 |
| *Dpp6* | 1.52E-09 | 2.29149917 | 3.16E-03 | 9.55E-01 | 1.00E+00 |
| *Top2a* | 1.51E-09 | 2.28346598 | 3.09E-03 | 9.56E-01 | 1.00E+00 |
| *Mypn* | 1.50E-09 | 2.28222579 | 3.08E-03 | 9.56E-01 | 1.00E+00 |

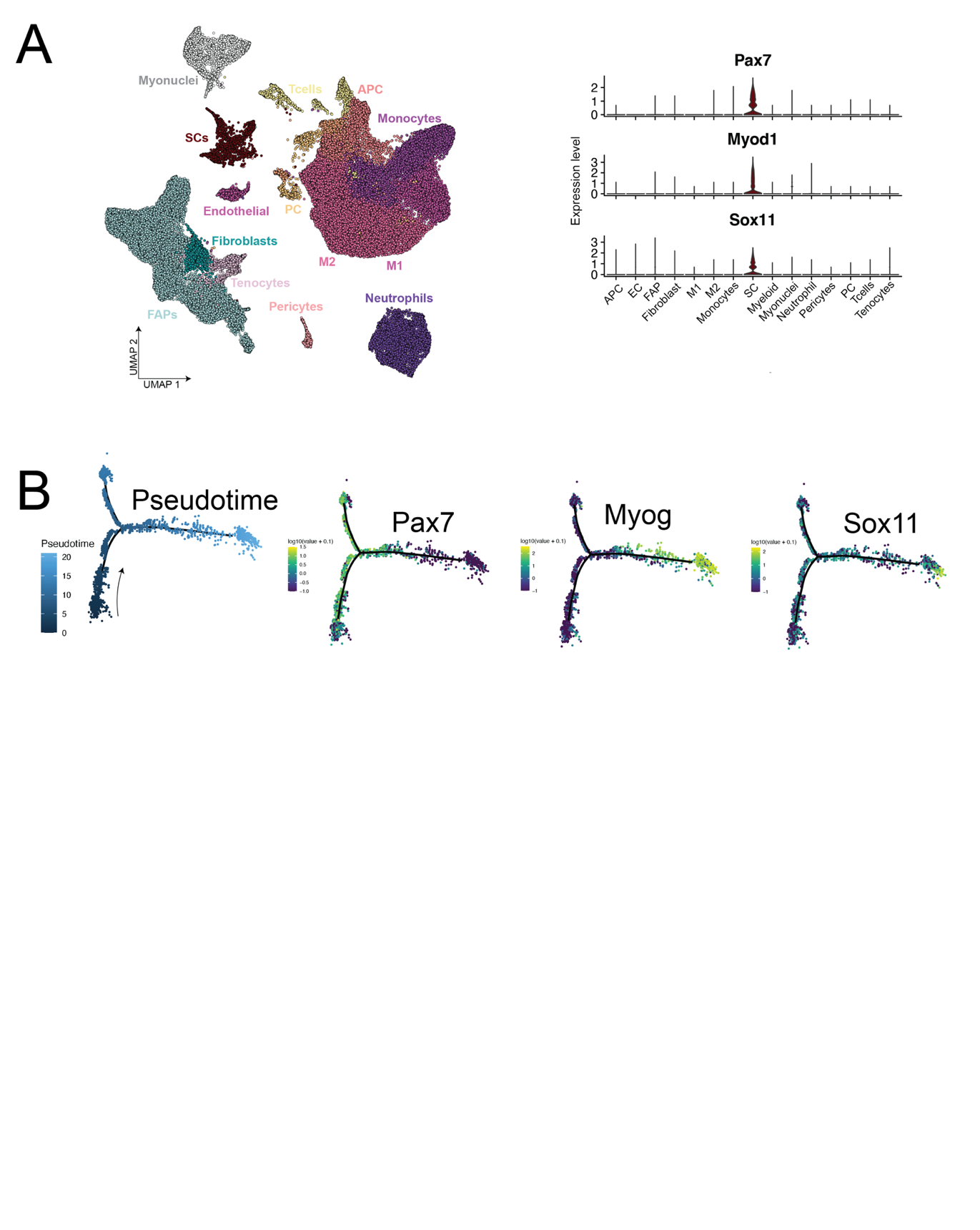

### Supplementary Figures

### Figure S1. Sox11 expression enriched in differentiating MuSCs. A. Left: UMAP of previously published scRNA-seq dataset on mononuclear cells from non-injured, 0.5, 2, 3.5, 5, 10, and 21 DPI aggregated together and labeled based on their broad cell type (right). Right: Violin plots to how the expression of *Pax7* and *Myod1*, which are specific to the MuSC subset, and *Sox11* whose expression is also specific to MuSCs. B. Monocle pseudotime trajectory analysis of MuSCs and gene expression plots on pseudotime of *Pax7*, *Myog* and *Sox11* from 5 DPI MuSCs shown in Fig. 1D. Related to Figure 1.

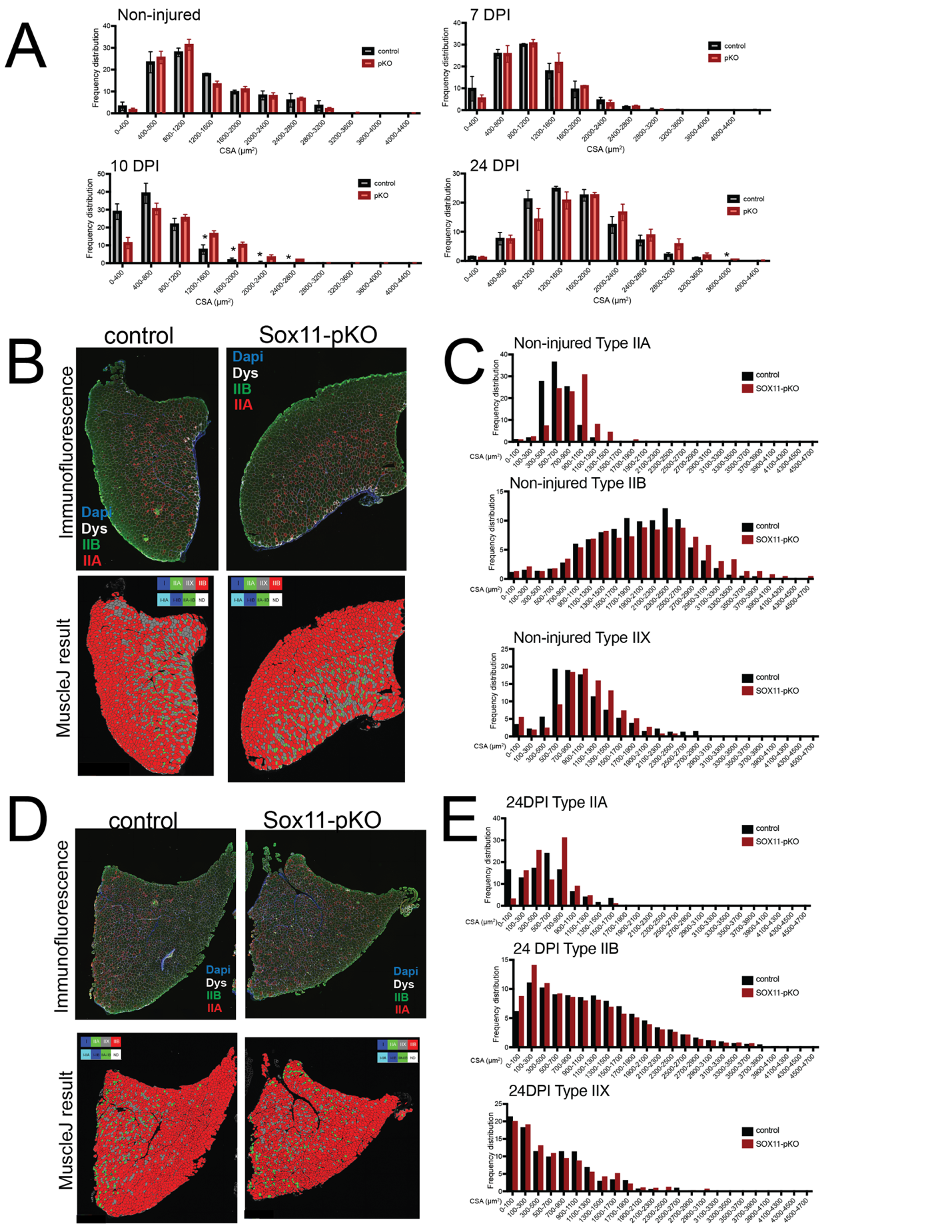

### Figure S2. Analysis of muscle fiber area and type for Sox11-pKO mice. A. Frequency distribution pots for CSA of TA muscle fibers from non-injured, 7, 10, and 24 DPI mice from control and Sox11-pKO mice. Measurements binned in 400 um^2^ bins. B. Representative immunofluorescence images to detect fiber Type IIA, Type IIB, Dystrophin and nuclei (DAPI) on non-injured muscles from control and Sox11-pKO mice (top panel), output from MuscleJ (bottom panel). C. Frequency distribution pots for CSA of TA muscle fibers from non-injured muscle, separated by inferred fiber type as analyzed from MuscleJ. Measurements binned in 400 um^2^ bins. D. Representative immunofluorescence images to detect fiber Type IIA, Type IIB, Dystrophin and nuclei (DAPI) on 24 DPI muscles from control and Sox11-pKO mice (top panel), output from MuscleJ (bottom panel). E. Frequency distribution pots for CSA of TA muscle fibers from 24 DPI muscle, separated by inferred fiber type as analyzed from MuscleJ. Measurements binned in 400 um^2^ bins. Related to Fig. 4.

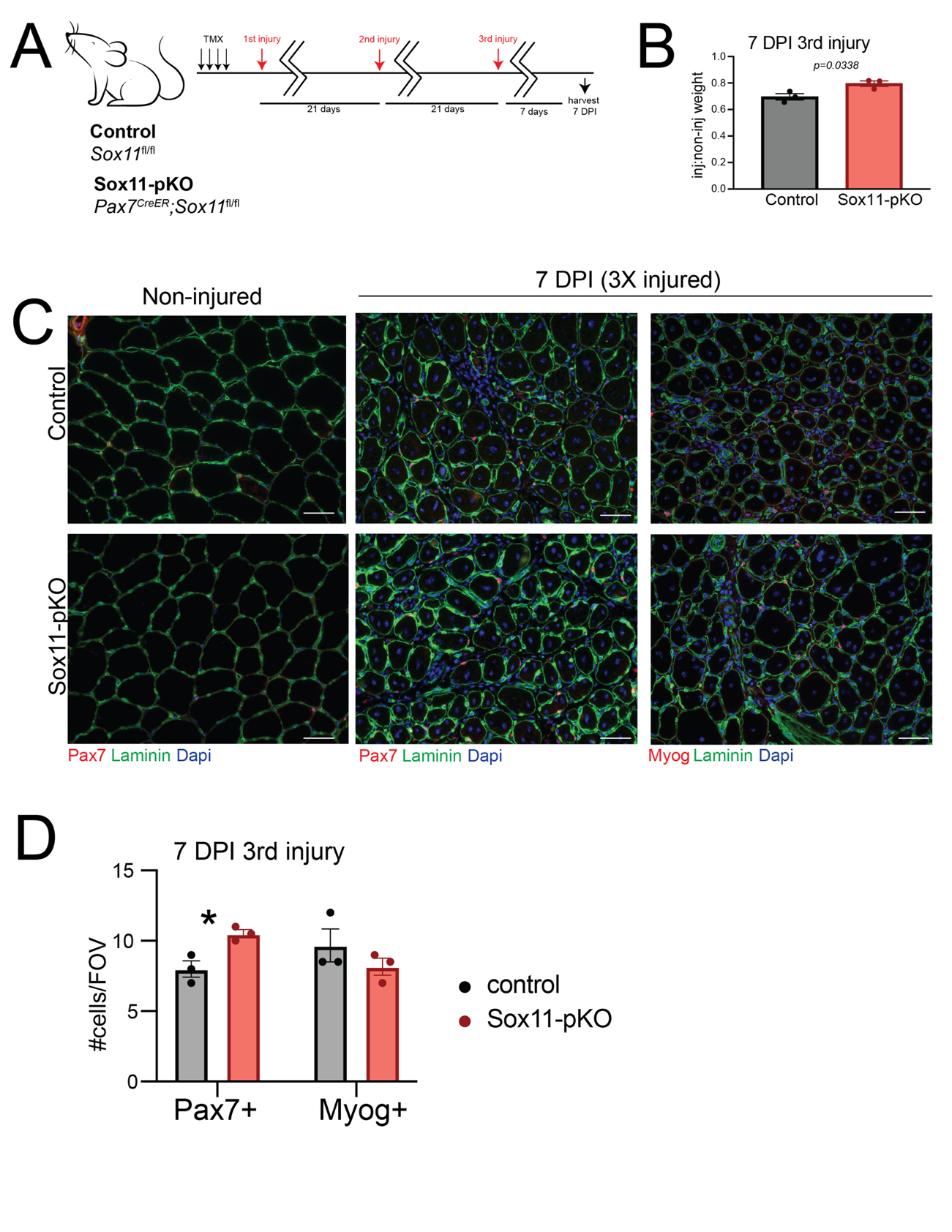

**Figure S3. Requirement of Sox11 for regenerative capacity of muscle stem cells after multiple rounds of injury. A.** Experimental outline to evaluate the requirement of Sox11 in MuSCs. Control and Sox11-pKO mice were injected with tamoxifen to induce recombination of the floxed allele. Subsequent MuSC function via muscle regeneration was evaluated using acute injury of the TA muscle with CTX 3 times (21 days between injury) and collected at 7 days post the 3^rd^ injury. **B.** Recovery TA weight, measured as the ratio of injured/non-injured contralateral muscle weight for 3X injured muscle. **C.** Representative images of immunofluorescence on TA muscle sections to detect Pax7 or Myog, Dystrophin, and nuclei (counterstained with DAPI) from control (top panel) and Sox11-pKO (bottom panel) mice at 0 (non-injured), 7 days post 3^rd^ injury **D.** Quantification of the number of Pax7+ cells/FOV and Myog+ cells/FOV for control and Sox11-pKO mice. Scale bars: 50 µm. *p-value = 0.01. Related to Figure 4.

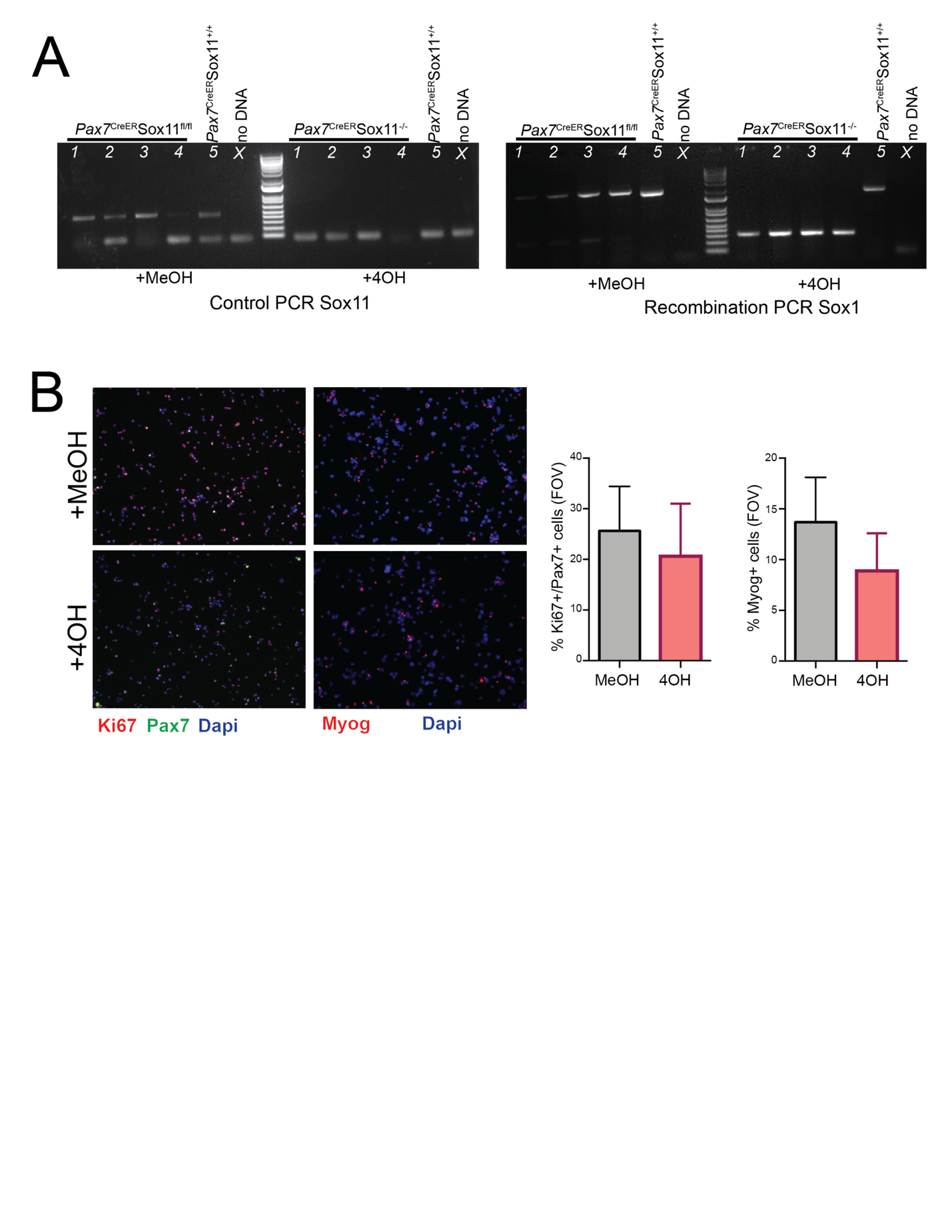

**Figure S4. In vitro analysis of Sox11-null myoblasts. A.** Recombination PCR to confirm the successful recombination of the Sox11-floxed allele upon the nuclear translocation of Cre recombinase, shown from myoblasts isolated from Sox11-pKO control mice treated with MeOH (top) or 4OH (bottom); protocol adapted from the Lefebvre lab. **B.** Primary myoblasts isolated from Sox11-pKO hindlimb muscle were cultured in the presence of MeOH (vehicle control) or 4-OH for 48 hours. Cells were seeded at equal concentrations, allowed to proliferate for 24 hours, fixed, and immunofluorescence was used to detect Ki67, Pax7, and nuclei (DAPI) (right panel) or Myog and nuclei (DAPI) (left panel). Bar graphs represent quantification of the number of Pax7+/Ki67+ cells per FOV or Myog+ cells per FOV. Related to Figure 5.

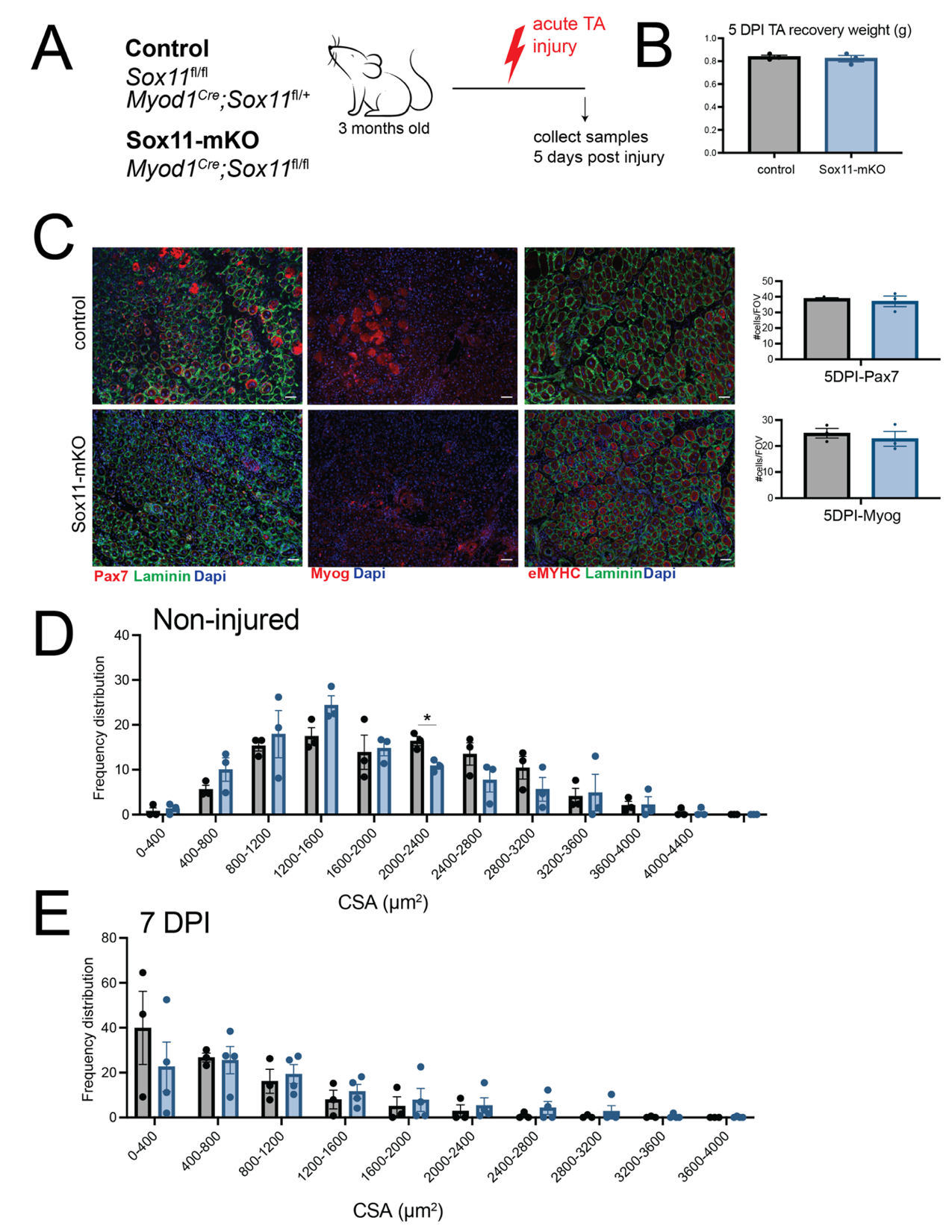

**Figure S5. Loss of Sox11 in muscle progenitors in adult and old mice. A.** Experimental outline to evaluate the requirement of Sox11 for muscle regeneration. Intramuscular TA injury was induced with CTX and samples were collected at 5 DPI. **B.** Recovery TA weight, measured as the ratio of injured/non-injured contralateral muscle weight for 5 DPI muscle. **C.** Representative images of immunofluorescence on TA muscle sections to detect Pax7 or Myog, Laminin and nuclei (counterstained with DAPI) from control (top panel) and Sox11-mKO (bottom panel) mice at 5 DPI and quantifications of the number of Pax7+ cells/FOV and Myog+ cells/FOV for control and Sox11-pKO mice based on 100X magnification FOV. Scale bars: 50 µm. Related to Figure 5.

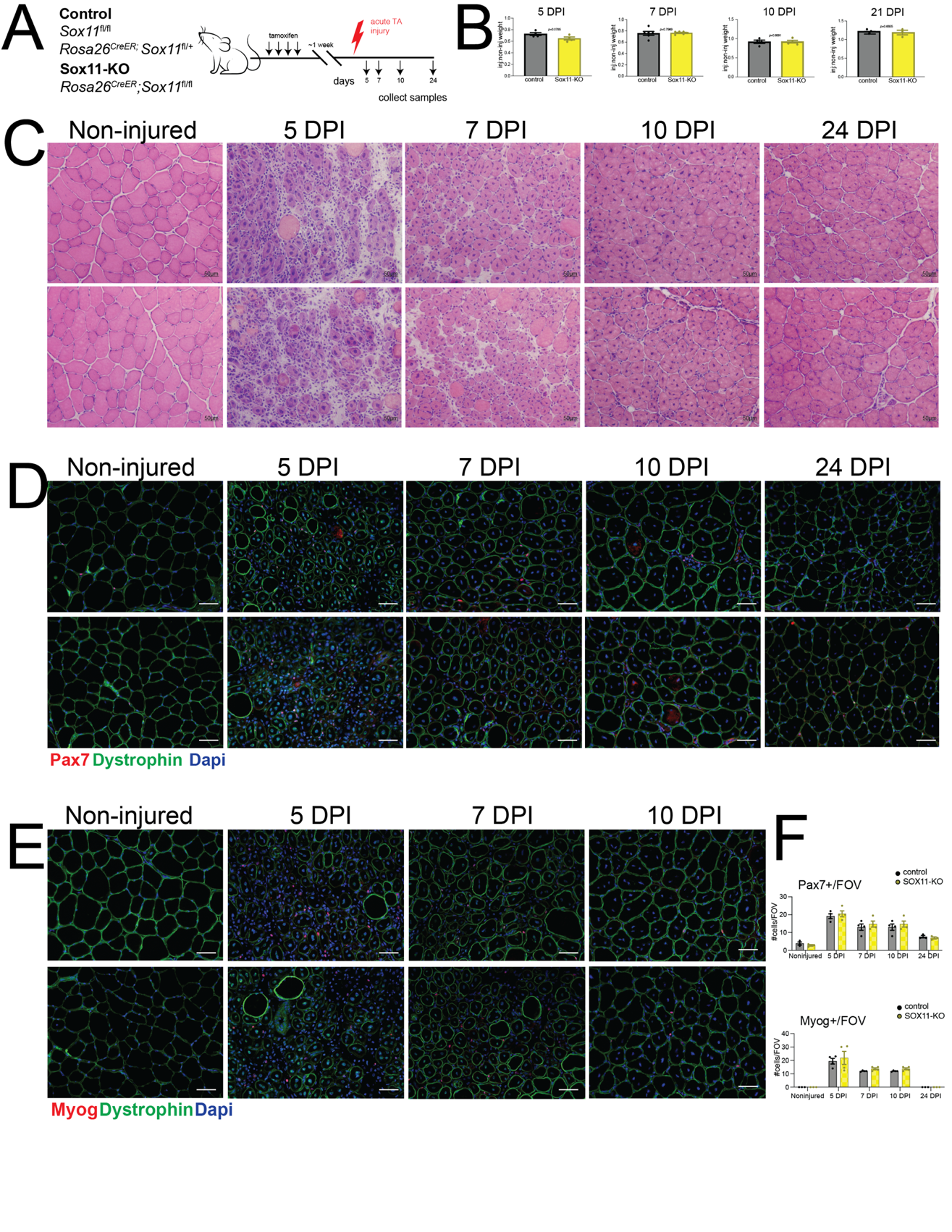

**Figure S6. Impact of global loss of Sox11 on muscle stem cell function and regeneration. A.** Schematic of tamoxifen injection, muscle injury with BaCl_2_ and the respective post-injury timepoints for muscle analysis. **B.** Recovery TA weight for control and Sox11-pKO mice, a measured by the ratio of injured: non-injured TA muscle. **C.** Representative images of H&E staining from cross-sections of non-injured, 7, 10 and 24 DPI TA muscle from control (top) and Sox11-KO (bottom). **D.** Representative immunofluorescence images of cross-sections from control (top) or Sox11-KO (bottom) stained with antibodies to detect Pax7, Dystrophin and nuclei (DAPI), of non-injured, 5, 7, 10 and 24 DPI muscle. **E.** Representative immunofluorescence images of cross-sections from control (top) or Sox11-KO (bottom) stained with antibodies to detect Myog, Dystrophin and nuclei (DAPI), of non-injured, 5, 7, and 10 DPI muscle. F. Quantification of the number of Pax7+ cells/FOV (top graph) and Myog+ cells/FOV (bottom graph) for control and Sox11-KO mice based on 200X FOV. Scale bars: 50 µm.
